# Dicer-mediated RNA silencing is the key regulatory mechanism in the biocontrol fungus *Clonostachys rosea*-wheat interactions

**DOI:** 10.1101/2023.09.24.559174

**Authors:** Edoardo Piombo, Ramesh Raju Vetukuri, Dan Funck Jensen, Magnus Karlsson, Mukesh Dubey

## Abstract

The intricate molecular interplay between beneficial fungi and plants is vital to plant growth promotion and induced defense response. This study explored the role of DCL-mediated RNA silencing in the interaction between the biocontrol fungus *Clonostachys rosea* and wheat roots. We investigated the impact of DCL (Dicer-like) gene deletions in *C. rosea* on its root colonization ability. Our results revealed that the deletion of *dcl2* significantly enhanced *C. rosea* biomass on wheat roots, indicating a pivotal role of DCL2 in root colonization. Transcriptome sequencing of *C. rosea* and wheat during their interactions unveiled extensive gene expression changes. In wheat, genes related to stress responses were upregulated during *C. rosea* interactions, while genes associated with plant cell wall modification and metabolic processes were downregulated, suggesting complex regulatory responses and a trade-off between defense mechanisms and growth promotion. Deletion of *C. rosea dcl1* and *dcl2* altered the transcriptomic responses of wheat roots during interactions. Wheat genes associated with stress responses were downregulated during interactions with DCL deletion strains. In contrast, genes involved in metabolic processes and growth were upregulated, emphasizing the cross-kingdom regulatory role of *C. rosea* small RNAs (sRNAs). We identified 18 wheat miRNAs responsive to *C. rosea* interactions. Furthermore, we identified 24 endogenous and six cross-kingdom potential gene targets for seven and five differentially expressed miRNAs, supported by their inverse gene expression pattern. In *C. rosea*, we found a large transcriptional reprogramming of genes during interaction with wheat roots. The upregulated genes were associated with carbohydrate and polysaccharide catabolic processes, membrane transporters and effectors. Conversely, downregulated genes were mainly associated with transition metal ion transport and homeostasis processes. The deletion of *dcl1* and *dcl2* had significant effects on gene expression. A higher number of genes upregulated in WT during the interaction were restored in DCL deletion mutants, suggesting DCL-mediated gene expression regulation. Furthermore, we identified 21 differentially expressed micro-RNA-like RNAs (milRNAs) in *C. rosea*; nine were DCL-dependent. They had putative gene targets in *C. rosea*, including transcription factors, effectors, transporters, and enzymes involved in specialized metabolite production. Cross-kingdom RNA silencing was also observed, with seven DCL-dependent *C. rosea* milRNAs potentially targeting 29 genes in wheat. These findings provide valuable insights into the molecular mechanisms underlying the beneficial interaction between fungi and plant roots. In addition, the study shed light on the role of sRNA-mediated gene regulation in the *C. rosea*-wheat interaction, with potential implications for sustainable agriculture and biocontrol strategies.

## Introduction

The genetic information flows from DNA to RNA to protein via transcription and translation, respectively. This flow of information is regulated at the transcriptional and post-transcriptional levels to maintain the proper functioning of cells. Post-transcriptional gene silencing (PTGS) is a highly conserved process of gene expression regulation, also called RNA silencing. This activity is performed by small non-coding RNAs (sRNAs) commonly ranging from 18-40 nucleotides (nt) in size (Hannon 2002; Ghildiyal and Zamore 2009; Huang et al. 2019). The sRNAs are typically produced from double-stranded RNAs and single-stranded RNAs with stem-loop by the enzymatic cleavage of endoribonuclease called Dicer (or Dicer-like [DCL] in fungi) producing small interfering RNAs (siRNAs) and microRNAs (miRNAs; microRNAs like RNAs in fungi [milRNAs]), respectively (Hannon 2002; Ghildiyal and Zamore 2009; Huang et al. 2019). Once produced, sRNAs are loaded onto an RNA-induced silencing complex (RISC) to guide Argonaute ribonucleases (AGO) to identify complementary messenger RNA (mRNA) for silencing through cleavage or modification of chromatin (Hannon 2002; Ghildiyal and Zamore 2009; Van Wolfswinkel and Ketting 2010). Due to their contribution to PTGS, sRNAs play a versatile role in living organisms’ life cycle, including biotic interactions (Rosa et al. 2018; Hudzik et al. 2020; Qiao et al. 2021). In addition, sRNAs can move bidirectionally and modulate communication between interacting organisms by regulating gene expression of recipient species through targeted gene silencing called cross-kingdom RNA silencing (Weiberg et al. 2013; Wang et al. 2016; Zhang et al. 2016; Cui et al. 2019; Hudzik et al. 2020; Wong-Bajracharya et al. 2022). Although the role of sRNAs in cross-kingdom RNA silencing between interacting organisms, including parasitic and mutualistic interactions, is established, their role in beneficial interactions between the fungal biocontrol agents and plant host is yet to be explored thoroughly.

Fungal biocontrol agents (BCAs), for example, species from the genera *Trichoderma* and *Clonostachys,* can occupy diverse ranges of environmental niches and closely interact at inter-species and intra-species levels for nutrients and space. In addition to directly antagonizing fungal plant pathogens, some of these species can colonize plant roots and establish mutual associations with host plants by promoting health and priming for the induced immune response against pathogens (Saraiva et al. 2015; Maillard et al. 2020; Tyśkiewicz et al. 2022). For successful beneficial association, biocontrol fungi and host plants reprogramme their genetic machinery and establish a molecular dialogue determining the degree of interactions (Zamioudis and Pieterse 2012; Lysøe et al. 2017; Macías-Rodríguez et al. 2020). RNA silencing has been shown to affect biocontrol fungi’s development, specialised metabolite production, and antagonistic activity (Carreras-Villaseñor et al. 2013; Cui et al. 2019; Piombo et al. 2021). Similarly, the role of fungal sRNA in biocontrol fungus-plant interaction is also considered. A recent study has shown that milRNAs of the biocontrol fungus *Trichoderma asperelleum* can target tomato genes involved in responses to ethylene and oxidative stress (Wang et al. 2021). Similarly, three wheat miRNAs engaged in response to abiotic and biotic stress are shown to be downregulated during the interaction with *Trichoderma cremeum* and *T. atroviride* (Salamon et al. 2021). However, such cross-kingdom action by *T. asperelleum* or wheat has not been experimentally proven. Furthermore, knowledge regarding how endogenous RNA silencing regulation could affect the relationship between biocontrol fungi and host plants is elusive. However, the role of plant sRNAs mediating the interaction between the biocontrol fungus *T. atroviride* and model plant *Arabidopsis thaliana* has recently been investigated (Rebolledo-Prudencio et al. 2022). In response to *T. atroviride* root colonization, *A. thaliana* showed induced expression of the RNA silencing machinery genes at local and systemic levels, which played a crucial role in the beneficial fungus-plant interactions by regulating expression pattern of the genes associated with plant growth and defense (Rebolledo-Prudencio et al. 2022). However, the precise plant sRNAs mediating the response to biocontrol fungi are unknown, as are their gene targets.

Our overall aim was to investigate sRNA-mediated mechanisms regulating interactions between biocontrol fungi and plant hosts to address whether sRNA-mediated interactions occur across a pathogenic-to-beneficial continuum of fungus-plant interactions, and thereby expand the understanding of cross-kingdom RNA silencing in fungus-plant interactions. We used the filamentous fungus *Clonostachys rosea* and wheat plant as the fungal BCA and plant host to achieve the goal. The fungus *C. rosea* can colonize plant roots including wheat and thereby promote plant health and induce an immune response (beneficial fungus-plant interactions) against several fungal plant pathogens (Sutton et al. 2002; Karlsson et al. 2015; Saraiva et al. 2015; Maillard et al. 2020; Dubey et al. 2014b, 2020). In addition, *C. rosea* can thrive as a necrotrophic mycoparasite and can antagonize plant pathogenic nematodes (Karlsson et al. 2015; Tzelepis et al. 2015; Iqbal et al. 2018; Sun et al. 2020; Broberg et al. 2021; Funck Jensen et al. 2021). To perform these functions, certain *C. rosea* strains are shown to regulate their genetic machinery and produce an arsenal of chemical compounds and proteins including hydrolytic enzymes, small secreted proteins and transporters (Dubey et al. 2014b, 2014a; Lysøe et al. 2017; Fatema et al. 2018; Nygren et al. 2018; Demissie et al. 2018, 2020; Broberg et al. 2021). The expression regulation of such compounds and proteins was recently shown to be mediated by sRNAs (Piombo et al., 2021, 2022). It is reported that deletion of *dcl2* resulted in mutants with altered expression of genes including those involved in the production of hydrolytic enzymes, membrane transporters, specialized metabolites and transcription factors (Piombo et al. 2021) ensuing reduced biocontrol ability of *C. rosea* against the fungal plant pathogen *F. graminearum* (Piombo et al. 2021).

The objectives of the current work are i) to identify *C. rosea* and wheat genes regulated during interactions, and ii) to unravel the role of sRNAs in mediating gene expression regulation during *C. rosea*-wheat interactions to understand the biocontrol mechanisms. We hypothesized that the transcriptomic response in *C. rosea* towards wheat plants is DCL-dependent. We further hypothesized that alteration in RNA silencing pathways in *C. rosea* will influence the transcriptomic response of wheat plants towards *C. rosea*. To achieve the objectives, we sequenced sRNAs and transcriptomes of *C. rosea* and wheat roots during interaction. We identified plant and fungal candidate miRNAs and genes with a potential role in biocontrol interactions. In addition, we used previously generated *dcl1* and *dcl2* gene deletion mutants (Piombo et al. 2021) to elucidate the role of sRNA on gene expression regulation at endogenous and cross-kingdom levels.

## Results

### Deletion of *dcl2* affects the root colonization ability of *C. rosea*

The role of DCL-mediated RNA silencing in beneficial fungus-plant interactions was investigated by comparing the root colonization ability of *C. rosea* WT and DCL deletion strains Δ*dcl1* and Δ*dcl2* (the *C. rosea* genome contains two genes *dcl1* and *dcl2* coding for DCL proteins) and the complementation strains (Δ*dcl1*+ and Δ*dcl2*+) generated in our previous study (Piombo et al., 2021). Root colonization ability was determined by measuring the biomass of *C*. *rosea* strains on wheat roots five (dpi) by quantifying the ratio between *C*. *rosea* DNA and wheat DNA with qPCR. Significantly (*P* < 0.019) higher *C*. *rosea*/wheat DNA ratios were detected on wheat roots inoculated with the Δ*dcl2* strain, compared with roots inoculated with the WT (**Figure 1**), suggesting increased biomass of the Δ*dcl2* strain, compared with the WT. Complementation strain Δ*dcl2*+ showed restoration of the phenotype. In contrast, no significant differences in *C*. *rosea*/wheat DNA ratio were found between WT and Δ*dcl1* inoculated wheat roots (**Figure 1**).

**Figure 1:**
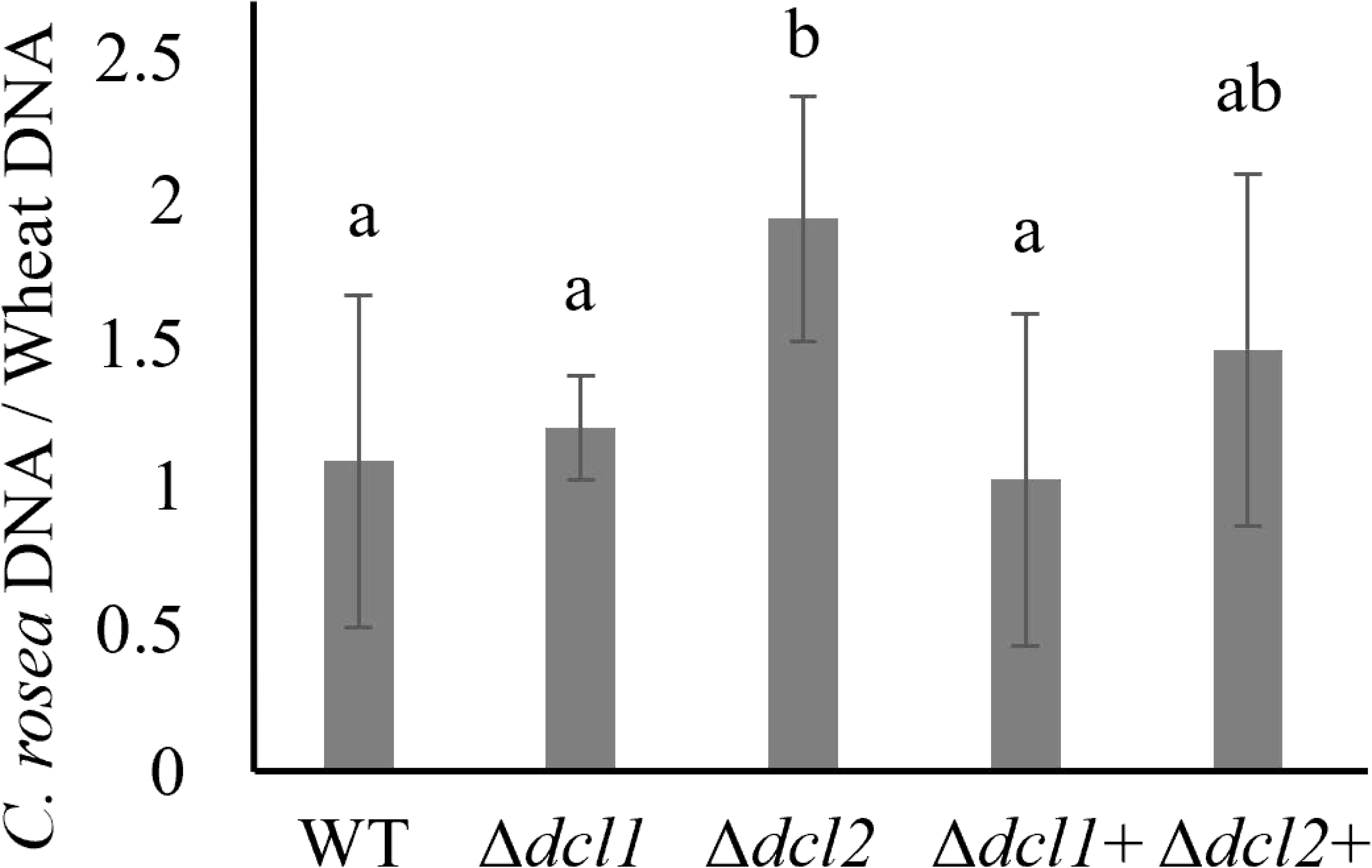
Determination of *C. rosea* root colonization in wheat roots, five dpi, by quantifying DNA level using RT-qPCR. *C. rosea* colonization is expressed as the ratio between *C. rosea* DNA and wheat DNA. Actin and Hor1 were used as target genes for DNA quantification for *C. rosea* and wheat, respectively.

### Transcriptome sequencing of *C. rosea* and wheat during interactions

To investigate transcriptomic (sRNAs and mRNAs expression) response of *C. rosea* strains and wheat during interactions, wheat roots (grown on moist filter plates in 9 cm Petri plates) inoculated with *C. rosea* conidia were harvested seven dpi and sRNA and mRNA expressions were analyzed by RNA-seq. Water inoculated wheat roots were used as a control for wheat transcriptome while *C. rosea* grown on moist filter paper was used as a control for *C. rosea* transcriptome. The mRNA sequencing produced between 33.51 and 26.03 million reads for each sample. Since the sequences contained read pairs from both the interacting species, the reads originating from *C. rosea* or wheat were identified by mapping to the respective genomes. As expected, in the samples coming from the interaction of *C. rosea* and wheat, the number of reads from *C. rosea* strains was significantly lower than the reads from wheat roots, amounting to 7% in the case of wildtype (WT) and 13% and 11% for the Δ*dcl1* and Δ*dcl2* mutants, respectively. A summary of the results from the mRNA sequencing is presented in **Table 1** and **Supplementary Table 1**.

**Table 1:** Number of mRNA reads assigned to wheat and *C. rosea*.

| Sample | *C. rosea* control | Wheat control | *C. rosea* – Wheat interactions |  |  |
| --- | --- | --- | --- | --- | --- |
| | | | WT-Wr | $\Delta dcl1$-Wr | $\Delta dcl2$-Wr |
| Raw reads (in million bp)<sup>a</sup> | 27.97 | 28.54 | 28.64 | 29.59 | 27.88 |
| Clean reads (in million bp)<sup>a</sup> | 27.83 | 28.29 | 28.35 | 29.36 | 27.73 |
| Reads unique to *C. rosea*<sup>#</sup> | 85.08 | 0.23 | 6.92 | 12.86 | 10.80 |
| Reads assigned to *C. rosea* genes<sup>#</sup> | 73.10 | NA | 5.74 | 10.95 | 9.37 |
| Unassigned *C. rosea* reads<sup>#</sup> | 11.12 | NA | 1.12 | 1.81 | 1.33 |
| Reads not mapped to *C. rosea*<sup>#</sup> | 0.85 | NA | 0.06 | 0.10 | 0.09 |
| Reads unique to wheat<sup>#</sup> | 0.55 | 85.44 | 76.10 | 70.36 | 74.02 |
| Reads assigned to wheat genes<sup>#</sup> | NA | 69.51 | 57.23 | 54.37 | 58.71 |
| Unassigned wheat reads<sup>#</sup> | NA | 13.39 | 11.54 | 10.01 | 10.32 |
| Reads not mapped to wheat<sup>#</sup> | NA | 2.54 | 7.32 | 5.98 | 5.00 |

The sRNA sequencing produced between 11.89 and 18.45 million reads per sample. After filtering out structural RNAs and reads shorter than 18 nt or longer than 32 nt, between 29% and 37% of reads were retained. Of these, between 21% and 30% of reads had an antisense match on the wheat genome, while between 2% and 6% of them on average mapped to the *C. rosea* genome. Conversely, up to 96% of these reads had a sense match on the wheat transcriptome, while this number never increased above 5% for *C. rosea* (**Table 2, Supplementary Table 1**).

**Table 2:**
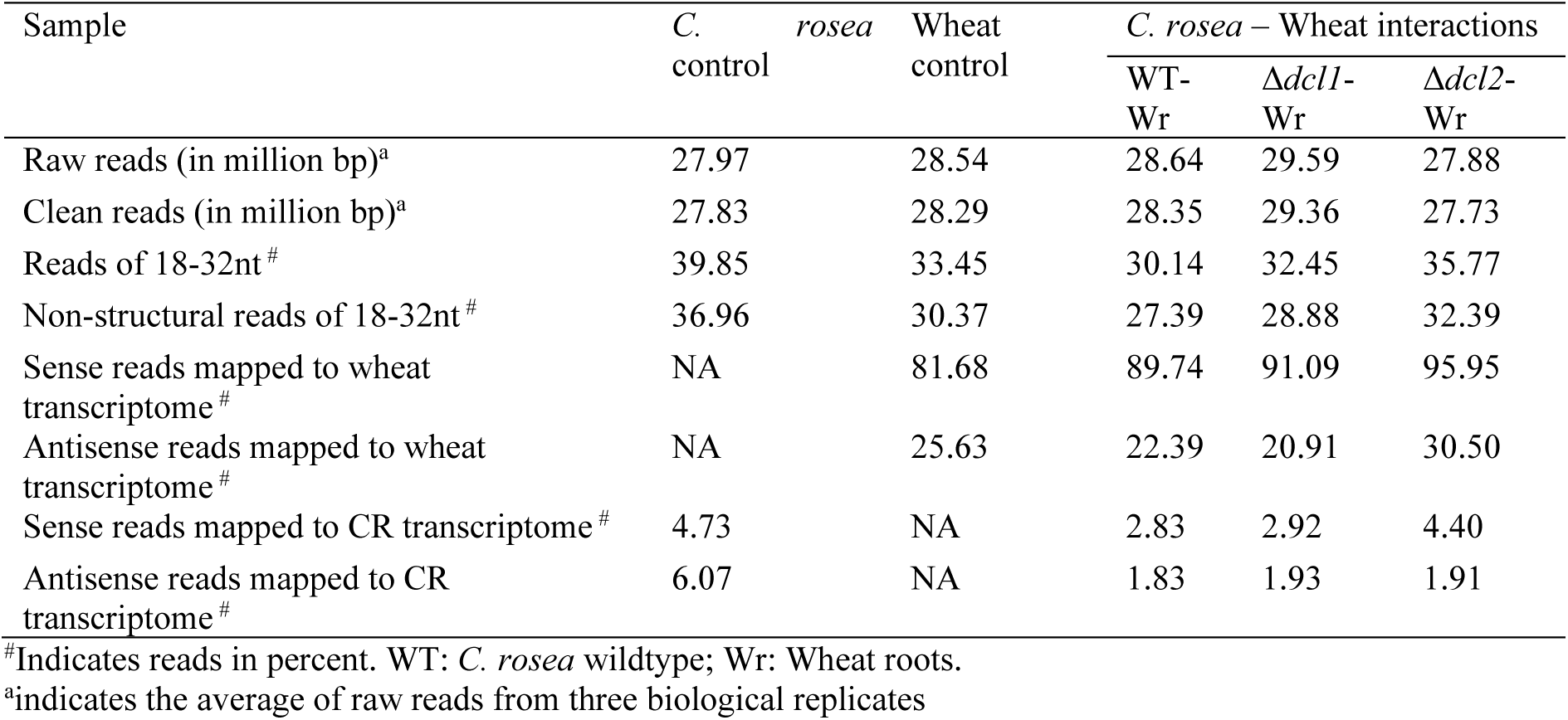
Number of sRNA reads assigned to wheat and *C. rosea*.

### The transcriptomic response of wheat during interaction with *C. rosea* involves genes associated with stress response and growth

To identify genes differentially expressed in wheat roots, the expression profile of wheat transcripts during interactions with *C. rosea* WT (Cr-Wr) was compared to that of non-interaction wheat control. In comparison to control, 280 wheat genes were significantly upregulated (log2FC > 1.5) and 208 were downregulated (log2FC < 1.5) in wheat during Cr-Wr (**Supplementary Table 2**). These genes were enriched in biological processes related to responses to several factors, including “response to acid chemical” (GO:0001101), “response to salt” (GO:1902074), “response to oxygen-containing compound” (GO:1901700) and “response to organic substance” (GO:0010033) (**Figure 2; Supplementary Table 3**). Among the upregulated genes 86 (31 % genes) were identified as biotic and abiotic stress related and wound-responsive. Fifty-three genes were associated with biotic stress tolerance consisting of LRR and lectin protein kinases, nodulin-like proteins, disease resistance proteins, defensins, vicilin-like proteins and ethylene responsive genes transcription factor. While abiotic stress-responsive genes consisted of late embryogenesis abundant (LEA) proteins (salt and oxidative stress tolerance) and dehydrins (dehydration and cold tolerance) (**Figure 3A, Supplementary Table 4).** The top 20 most upregulated genes include a disease resistance protein RGA5-like; *Fusarium* resistance orphan protein Traescs4b01g106100.1; defensin-like 1 protein, which has antifungal activity (Yan et al. 2015); vicilin-like seed storage proteins, known for inhibiting the spore germination and growth of filamentous fungi (Chung et al. 1997; Gomes et al. 1998); a dehydrin, involved in response to dehydration and cold (Shakirova et al. 2016). In this group we also identified a ubiquitinyl hydrolase 1 and an F-box protein, which affect several plant processes including stress response (Stefanowicz et al. 2015), as well as a protein TAR1-like, putatively involved in auxin biosynthesis and consequently in hormone crosstalk and plant development (Stepanova et al. 2008; Čarná et al. 2014) (**Supplementary table 5)**.

**Figure 2:**
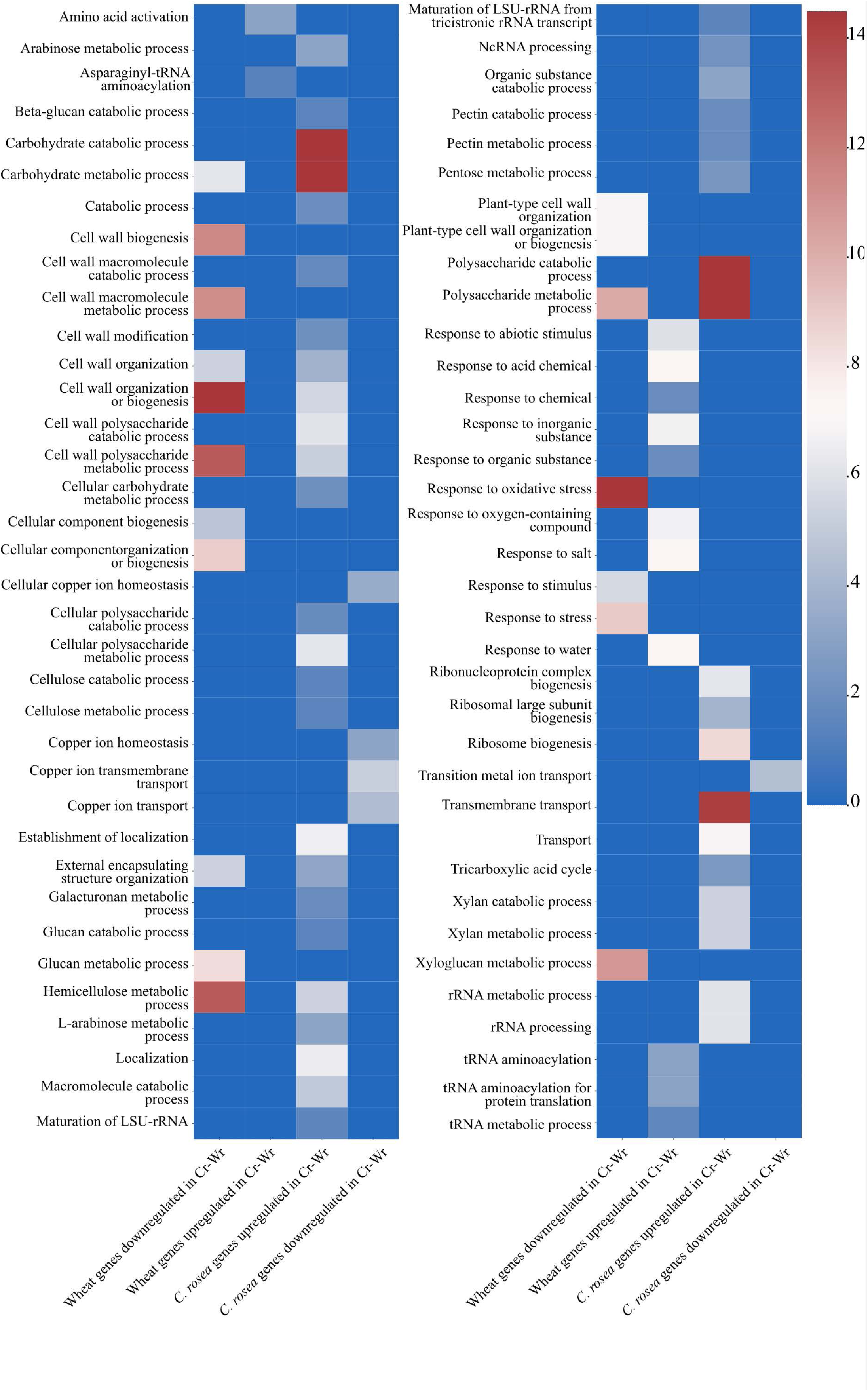
Gene ontology terms referring to biological processes enriched in wheat genes or *C. rosea* genes differentially expressed during the interaction between the two organisms.

**Figure 3:**
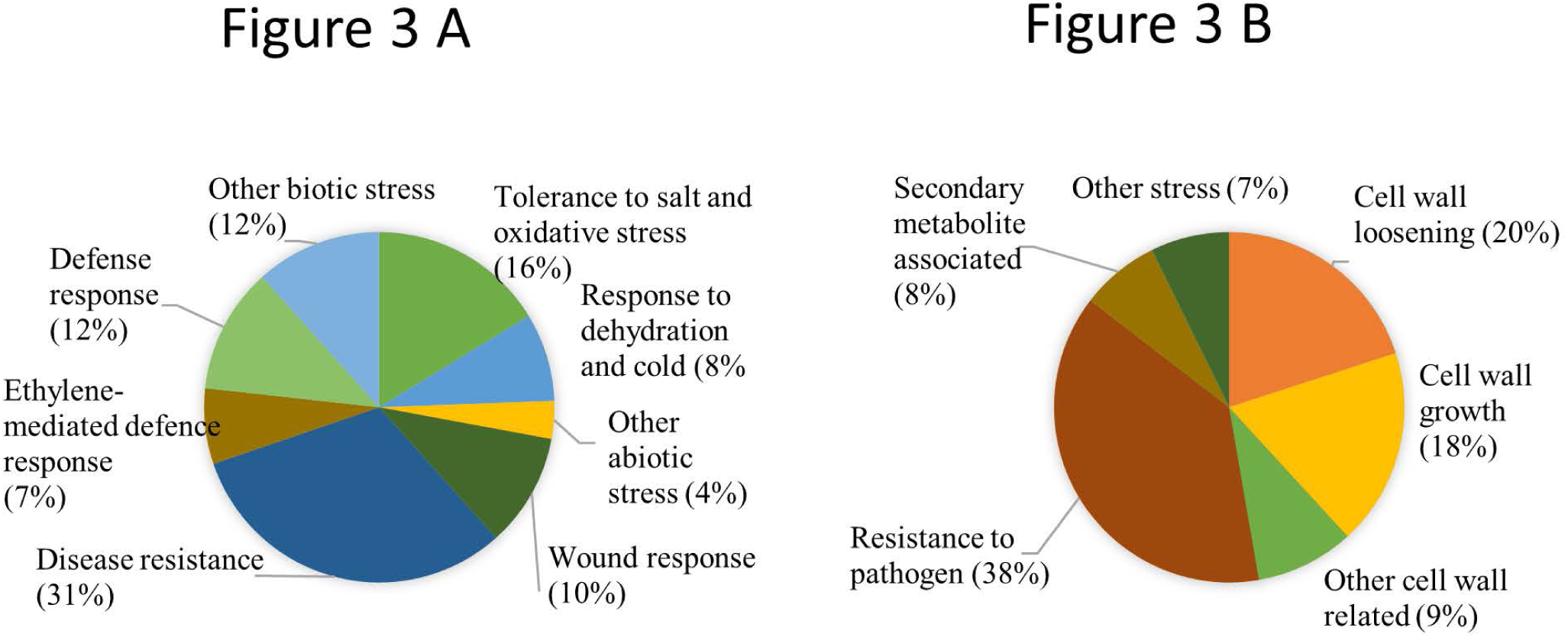
**A:** Pie chart showing **the** proportion of the stress-related genes in wheat upregulated during Cr-Wr. **B**: **Pie chart showing the** proportion of stress-related genes, secondary metabolism, and cell wall-related genes downregulated during Cr-Wr.

During the Cr-Wr 208 wheat genes were downregulated (**Supplemental Table 2**). These genes were enriched in a high number of biological processes, but mostly related to modification of plant cell wall and metabolic processes (**Figure 2**). The terms “response to oxidative stress” (GO:0006979), “response to stress” (GO:0050896), and “response to stimulus” (GO:0050896) were also enriched (**Figure 2, Supplementary table 3**). Among the downregulated genes, 55 (26 %) were associated with resistance to pathogens, and cellular growth by performing cell wall loosening and modification (**Figure 3B, Supplementary Table 4).** The downregulated pathogen responsive genes consisted of peroxidases associated with reactive oxygen burst, disease resistance proteins, pathogenesis-related proteins, and zealexin synthase and chalcone synthase involved in phytoalexin and flavonoid biosynthesis (**Figure 3B, Supplementary Table 4)**. The 26 downregulated genes consisting of genes coding expensing-like proteins and xyloglucan endotransglucosylase/hydrolase, polygalacturonase, cellulose synthase-like protein D1 and dirigent protein 5-like proteins predicted to be involved in cell wall loosening and modification (**Figure 3B, Supplementary Table 4)**. The list of top 20 significantly down-regulated genes include genes associated with plant defence against biotic and abiotic stresses. The identified genes were a NBS-LRR resistance proteins mediating pathogen sensing and host defense (DeYoung and Innes 2006; Sekhwal et al. 2015); a pathogenesis-related protein 1 with antifungal activity (Breen et al. 2017); a peroxidase 5-like involved in ROS burst; a zealexin A1 synthase-like involved in the synthesis of protective phytoalexins (Shen et al. 2019); two chalcone synthases, which affect resistance by influencing the salycilic acid response and the accumulation of flavonoid phytoalexins (Dao et al. 2011; Jayaraman et al. 2021); and a serpin, a class of protease inhibitors that can have a role in the inhibition of plant hypersensitive responses (Lema Asqui et al. 2018) (**Supplementary Table 5)**. These results suggest that the root colonization by *C. rosea* resulted in transcriptional reprograming of wheat gens associated with stress-response and growth indicating plausible trade-off between defense and growth.

### *Clonostachys rosea* interactions with wheat roots triggered transcriptional reprograming of gens coding for CAZymes, membrane transporters effector

In *C*. *rosea*, 1908 genes were upregulated during Cr-Wr, compared to *C*. *rosea* control, while 1262 were upregulated (**Supplementary Table 2**). The biological processes in the genes upregulated during Cr-Wr interactions mostly referred to an increase in the catabolism in many types of carbohydrates, and they included “carbohydrate catabolic process” (GO:0016052) and “polysaccharide catabolic process” (GO:0000272). This can be attributed to the fact that 229 of the upregulated genes were coding for putative CAZymes, and the majority of them (163) was reported as secreted in previous study (Piombo et al. 2023). Among the CAZymes, 134 were glycoside hydrolases, and 87 of them had CBM domain, the most frequent of which was CBM1, present in 55 proteins Even if glycoside hydrolases were the most common type of CAZyme, the most numerous single class was AA9 (21 genes), involved in the degradation of cellulose. CAZymes are also involved in cell wall organization and modification, whose GO terms are enriched as well in the upregulated genes (GO:0071554, GO:0042545). The remaining terms suggest a modification in rRNA metabolism and transport, and they include “ribosome biogenesis” (GO:0042254), “rRNA metabolic process” (GO:0016072), “localization” (GO:0051179) and “transmembrane transport” (GO:0055085) (**Figure 2**). The transmembrane transport, in particular, can be related to the upregulation of 135 MFS and 13 ABC transporters, classes that have been proven important in the antagonistic activity of *Clonostachys* species (Broberg et al. 2021). In particular, the most numerous classes of upregulated MFS transporters are 2.A.1.1(sugar porters), 2.A.1.14 (organic cation transporters) and 2.A.1.2 (drug: H^+^ antiporter), with 45, 31 and 28 members respectively. The enriched cellular component GO terms are “extracellular region” (GO:0005576), “preribosome” (GO:0030684) and “nucleolus” (GO:0005730), suggesting these are the destination of the transmembrane transport. In addition, we identified 34 upregulated effector genes during Cr-Wr compared to the control (**Supplementary Table 3**). The 20 *C. rosea* genes most upregulated during Cr-Wr encoded nine proteins with roles in the degradation of the plant cell wall, in the form of three monoxygenases of the AA9 class, two GH12 cellulases, one alpha-L-arabinofuranosidase GH54 and one acetyl xylan esterase CE1, both involved in hemicellulose degradation and a glucuronyl esterase CE15 which hydrolyzes lignin-carbohydrate ester linkages between lignin and glucuronoxylan (Mosbech et al. 2018; d’Errico et al. 2016; Arnling Bååth et al. 2016). Moreover, 13 of these 20 genes encode for putative secreted effectors, suggesting some role in affecting the plant immunity response (**Supplementary Table 6)**.

The GO term enrichment analysis of downregulated genes highlighted only 4 biological processes associated with “transition metal ion transport” (GO:0000041), “copper ion transport” (GO:0006825), “copper ion transmembrane transport” (GO:0035434), and “cellular copper ion homeostasis” (GO:0006878) (**Figure 2, Supplementary table 3**). The 20 *C. rosea* genes most downregulated during the interaction with wheat included 3 transcription factors, a secreted putative effector, a NRPS involved in the synthesis of an unknown specialized metabolite, a superoxide dismutase participating in defense from reactive oxygen species (ROS) and four membrane transporters, three of which were predicted to mediate the transport of copper, iron and zinc (**Supplementary Table 6)**. Taken together, these data highlight that the *C. rosea* interactions of wheat roots triggers the transcriptional reprograming of genes associated with stress-response, catabolism, and transport.

### Deletion of *C. rosea dcl1* and *dcl2* altered the transcriptomic response of wheat root during the interactions

To investigate whether deletion of *dcl1* and *dcl2* in *C. rosea* can affect the gene expression regulation in wheat root during Cr-Wr, the gene expression pattern of wheat roots during the interaction with DCL1 deletion strain (Δ*dcl1*-Wr) and DCL2 deletion strain (Δ*dcl2*-Wr) was analyzed and compared to Cr-Wr. We identified one 144 wheat genes commonly upregulated during the interaction with the DCL deletion mutants compared to wheat control, but not during Cr-Wr. While 93 and 78 were upregulated only during Δ*dcl1*-Wr and Δ*dcl2*-Wr, respectively (**Figure 4**). On the contrary, only eleven genes were downregulated during the response to both mutants but not to the WT, while 114 and 119 were uniquely downregulated during Δ*dcl1*-Wr and Δ*dcl2*-Wr, respectively. The differentially expressed wheat genes could be divided into nine modules, which showed how the deletion of *C. rosea dcl1* had a lesser effect on wheat response than the deletion of *dcl2*, as every module differed in expression between Cr-Wr and Δ*dcl2*-Wr, while ME_1 and ME_2 were minimally affected by the deletion of *dcl1* (**Supplementary Figure 1**). In summary, our result showed a shift in the transcriptomic response of wheat roots during Δ*dcl1*-Wr or Δ*dcl2*-Wr suggest a potential role of *C. rosea* sRNAs in Cross-kingdom RNAi.

**Figure 4:**
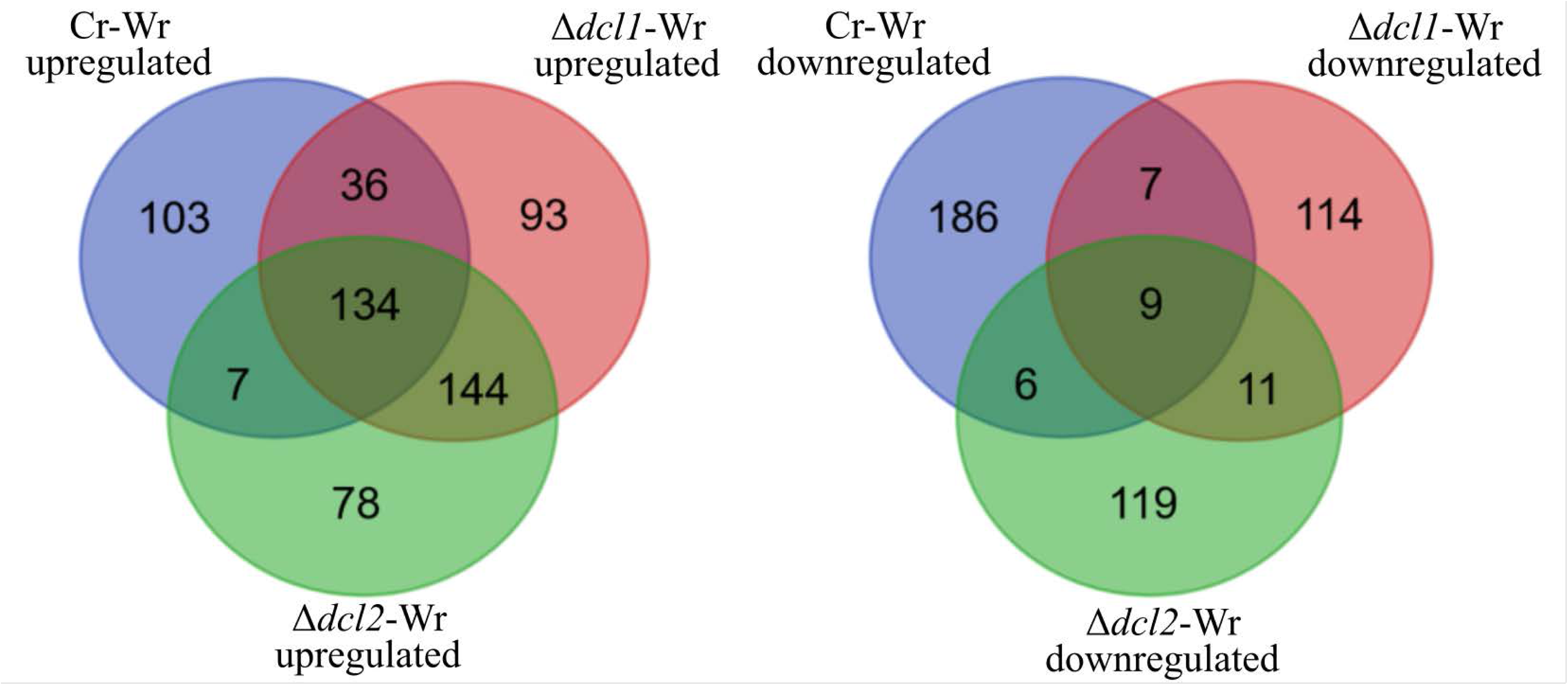
Number of differentially expressed wheat genes during the interactions with *C. rosea* WT or DCL gene deletion strains. Venn diagram generated with https://bioinformatics.psb.ugent.be/webtools/Venn/

### Wheat interaction with *dcl* deletion strains affects expression pattern of genes associated with stress response, metabolism and growth

Among 146 wheat genes, which were upregulated during Cr-Wr but not during either Δ*dcl1*-Wr or Δ*dcl2*-Wr, 65 were associated with stress response (**Figure 5, Supplementary Table 7**). The Go term analysis showed the all terms enriched in these genes were related to the response to several stress-related factors (**Figure 6A**). More specifically, this group included 2 protein phosphatases 2C, interacting with ARF genes involved in resistance to powdery mildews and abiotic stresses (Li et al. 2021), as well as 11 LEA proteins, necessary for tolerance of salt and oxidative stress (Koubaa and Brini 2020). Both of these protein classes are regulated by abscisic acid, and the same is true for membrane proteins PM19L-like, one of which had DCL-dependent expression in the current study (Chen et al. 2015; Qi et al. 2016; Nguyen et al. 2019).

**Figure 5:**
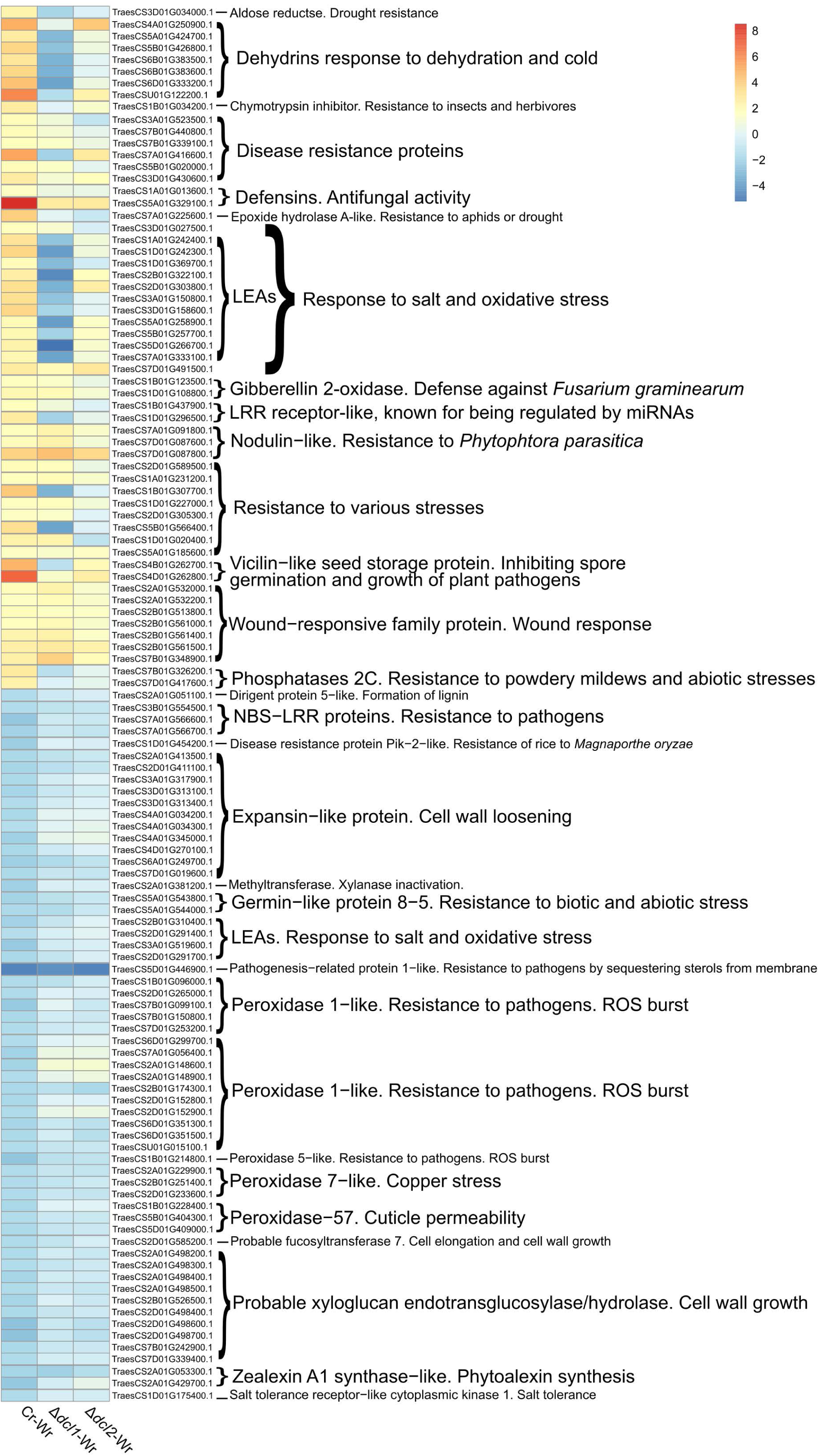
The heatmap shows the expression (Log2FC) of selected wheat genes of interest. All genes were differentially expressed during Cr-Wr but during both or either Δ*dcl1*-Wr or Δ*dcl2*- Wr.

**Figure 6:**
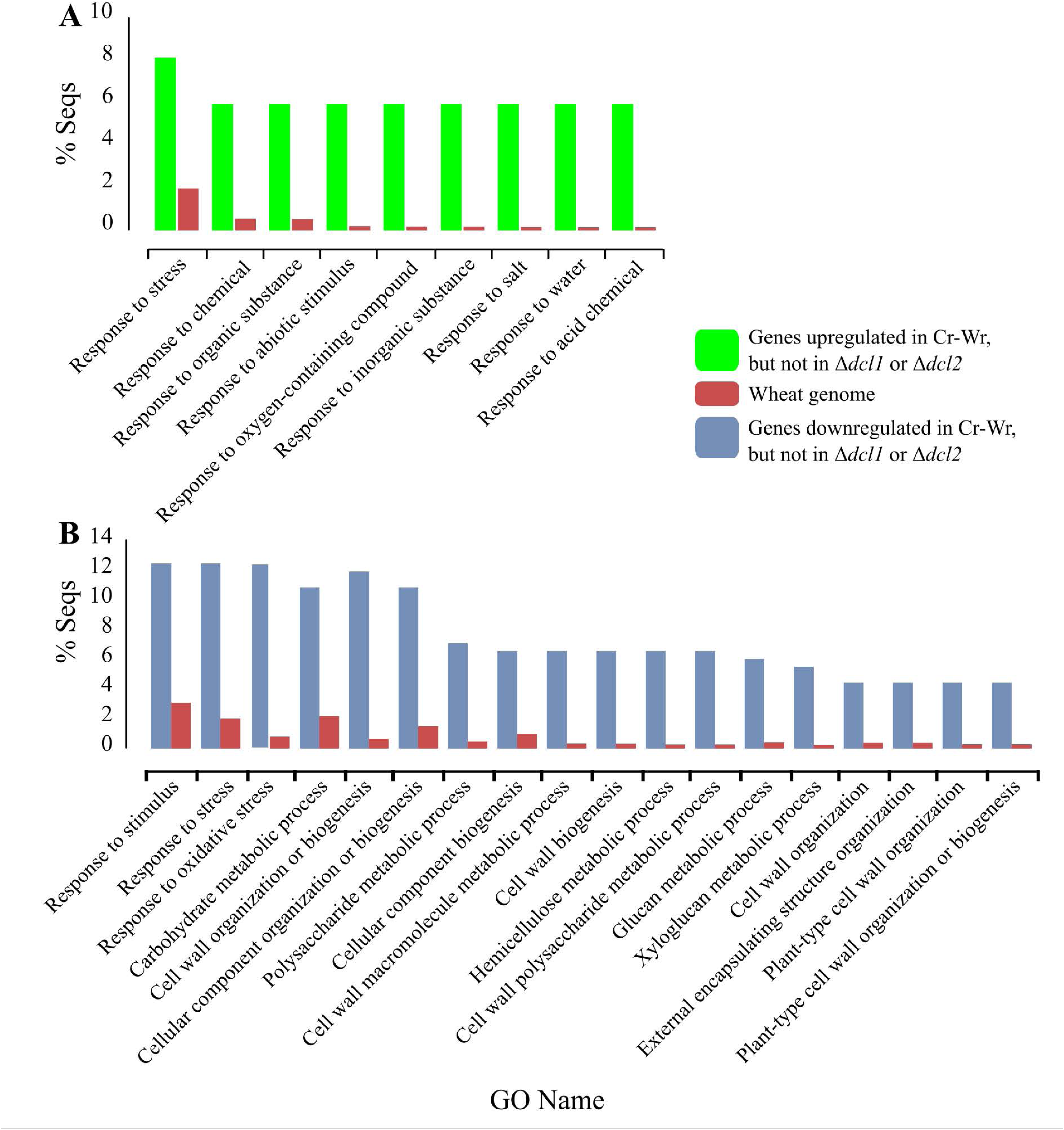
Gene ontology terms referring to biological processes enriched in wheat genes or *C. rosea* genes differentially expressed during the interaction between wheat and the WT, but not when the plant interacted with the Δ*dcl1* or Δ*dcl2* mutant.

Many other DCL-responsive genes are related to resistance to various types of abiotic stresses. We could detect seven DHN dehydrins, involved in the response to dehydration and cold (Shakirova et al. 2016), one H-type thioredoxin, mediating responses to oxidative stresses (Wu et al. 2017; Shaw et al. 2017) and one aldose reductase, whose overexpression improves drought resistance in transgenic plants (Fehér-Juhász et al. 2014). Other detected genes were involved in resistance to pathogens, such as one Bowman-Birk type trypsin inhibitor-like isoform X2, which can inhibit *in vitro* growth of *F. graminearum*, *Fusarium culmorum* and *Fusarium tritici* (Chilosi et al. 2000), and one premnaspirodiene oxygenase-like protein, involved in resistance to *Phytophthora capsici* in black pepper (Paul et al. 2019). Two defensin proteins also fall in this group, and similar proteins have an antifungal activity carried out through cell wall permeabilization (Yan et al. 2015). We also identified two vicilin-like seed storage proteins, a class known for inhibiting spore germination and growth of several filamentous fungi (Chung et al. 1997; Gomes et al. 1998), and two LRR receptor-like serine/threonine-protein kinase, a class involved in resistance to *Puccinia triticina* and *Plasmopara viticola*, and also known for being regulated by miRNAs (Zhao et al. 2015; Fu et al. 2020; Lee et al. 2020). Other DCL-dependent genes seem to be involved in resistance to both biotic and abiotic stresses, such as an epoxide hydrolase A-like, similar to genes involved in resistance to aphids and others targeted by drought-responsive miRNAs (Hua et al. 2019; Tulpová et al. 2019), or one NAC protein, a class involved in drought resistance, sensitivity to ABA, lignin biosynthesis and resistance to *F. graminearum* and *Puccinia triticina* (Mergby et al. 2021; Soni et al. 2021; Zhang et al. 2021) (**Figure 5: Supplementary Table 7**). In summary, this data suggests that the interaction of wheat with *C. rosea* DCL deletion strains resulted in the downregulation of stress-responsive genes.

Deletion of DCL genes restored the expression of 199 wheat genes during Δ*dcl1*-Wr or Δ*dcl2*- Wr, which were downregulated during Cr-Wr (**Supplementary Table 2, Figure 4**). These genes were enriched mainly in biological processes related to the synthesis, organization and modification of the cell wall, including for example “plant-type cell wall organization” (GO:0009664), “external encapsulating structure organization” (GO:0045229), “hemicellulose metabolic process” (GO:0010410) and “xyloglucan metabolic process” (GO:0010411) (**Figure 6B, Supplementary Table 3**). Eleven of these genes are coding for expansin-like proteins with a role in plant cell wall loosening, while three others are peroxidases-57, whose overexpression increases cuticle permeability in *Arabidopsis thaliana* (Survila et al. 2016). The terms “response to oxidative stress” (GO:0006979) and “response to stress” (GO:0006950) were also enriched in this group of genes (**Figure 6B**), possibly due to the presence of several transcripts with roles in disease resistance. These are a methyltransferase involved in xylanase inactivation during *F. graminearum* infection (Tundo et al. 2015), two biosynthesis enzymes of phytoalexin zealexin A1 (Shen et al. 2019), three resistance proteins of class NBS-LRR, 14 peroxidases of classes 1, 2, and 5, involved in resistance to biotic stress (Choi and Hwang 2012; Peters et al. 2017; Khaleghi et al. 2021), and a disease resistance protein Pik-2-like, involved in resistance of rice to *Magnaporthe oryzae* and upregulated in wheat during *Blumeria graminis* infection (Nie and Ji 2019; Varden et al. 2019) (**Figure 5; Supplementary Table 7**). In summary, this data suggests that deletion of *dcl* in *C. rosea* resulted in the upregulation of wheat genes associated with metabolic processes, growth, and stress response.

### DCLs-mediated gene expression regulation in *C. rosea* during interaction with wheat roots

The deletion of the *dcls* affected the expression of multiple *C. rosea* genes during the interaction with wheat roots. Five hundred and twelve genes were upregulated in the Δ*dcl1* mutant but not in the WT, while this number was 431 for the Δ*dcl2* mutant. The number of down-regulated genes in the mutants but not in the WT corresponded to 591 genes in Δdcl1 and 684 in Δ*dcl2* (**Figure 7**). The differentially expressed *C. rosea* genes could be divided into nine modules, each showing a significant difference in expression between the WT and the deletion mutants. The only exceptions were ME_1, similar in WT and Δ*dcl1,* and ME_5, identical in WT and Δ*dcl2* (**Supplementary Figure 2**).

**Figure 7:**
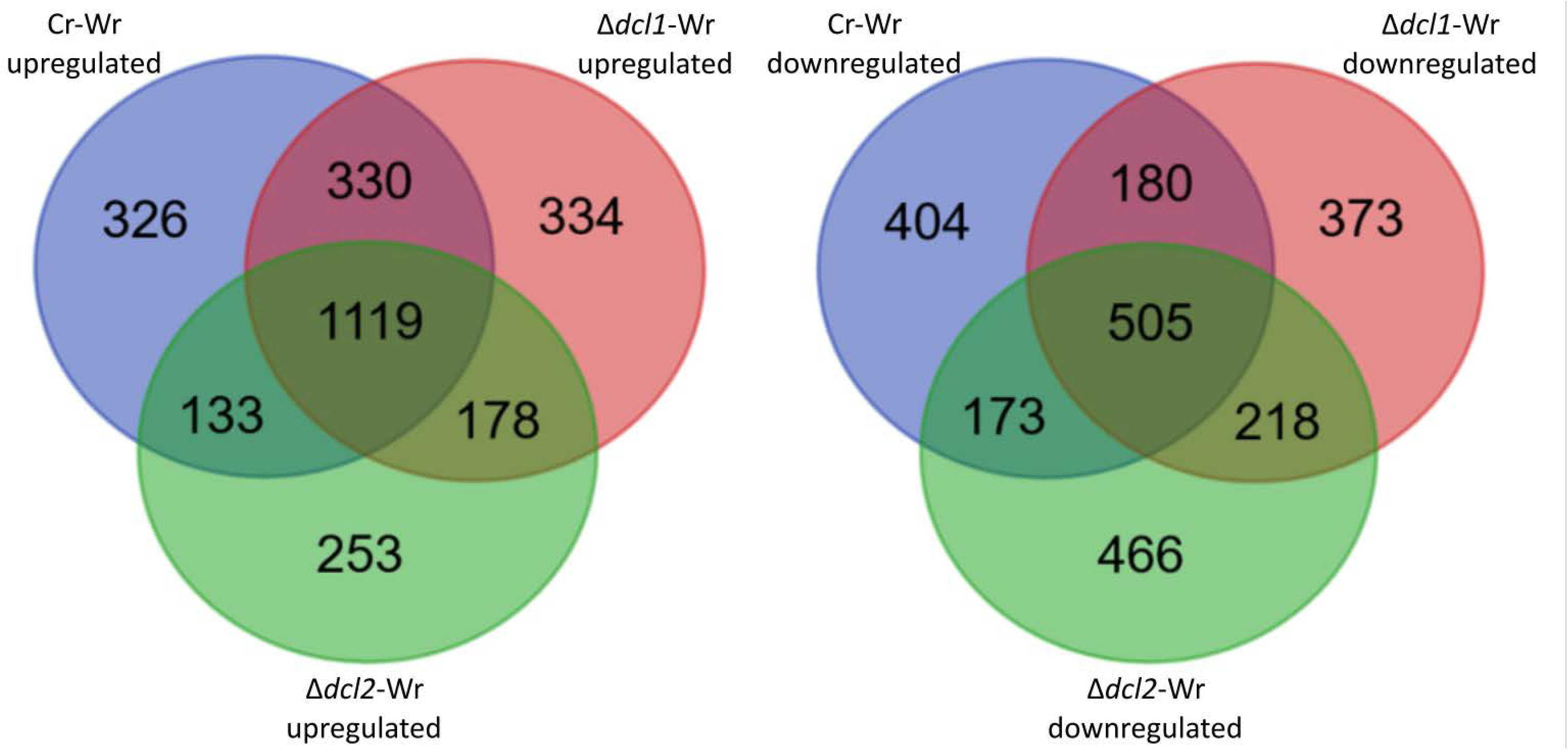
Number of differentially expressed *C. rosea* genes during the interaction with wheat roots. Venn diagram generated with https://bioinformatics.psb.ugent.be/webtools/Venn/

Similarly, the expression of 789 genes, which was upregulated during Cr-Wr, was restored during Δ*dcl1*-Wr or Δ*dcl2*-Wr, indicating DCL-mediated gene expression regulation. These genes were enriched in GO-term biological processes related to rRNA production, such as “ribosome biogenesis” (GO:0042254), “maturation of 5.8S rRNA” (GO:0000460), “maturation of LSU-rRNA” (GO:0000470) and “rRNA processing” (GO:0006364) (**Supplementary Table 3**).

Since many genes coding for CAZymes and effectors were upregulated during Cr-Wr, we wanted to know if their expression is DCL-mediated. The expression of 54 CAZymes genes was significantly downregulated in *C. rosea* DCL deletion strains during the interaction with wheat roots compared to the WT, 32 of which were predicted to be secreted in a previous work (Piombo et al., 2023). (**Supplementary table 7**). A higher number of genes (49 genes) were downregulated during Δ*dcl2*-Wr compared to the number of genes (25 genes) during Δ*dcl1*- Wr. The expression of 34 putative effector coding genes was downregulated in C. rosea dcl strains compared to the WT. Twenty-seven and 23 genes were downregulated during Δ*dcl2*- Wr and Δ*dcl1*-Wr, respectively, compared to Cr-Wr (Supplementary Table S7). Conversely, 757 genes, which were downregulated during Cr-Wr but not during Δ*dcl1*-Wr or Δ*dcl2*-Wr, were not enriched in any GO term.

### Identification of wheat miRNAs responsive to *C. rosea* interactions and their putative endogenous and cross-kingdom gene targets

To provide insights into gene expression regulation during Cr-Wr, we investigated sRNA-mediated wheat gene expression regulation by analyzing sRNA characteristics and their expression patterns in response to *C. rosea* root colonization. The length distribution of sRNA reads showed a higher proportion of reads with 20 nt (34%) and 24 nt of length (**Figure 8A**). The analysis of the 5’ terminal nucleotide composition showed a higher proportion (47-52%) of the reads with 5’ end adenine (5’ - A). At the same time, both guanine and uracil were present at the 5’ of 20% of the reads and cytosine was the lowest base in that position, covering less than 10% of the reads (**Figure 8B**). We compared the characteristics of wheat sRNAs produced in the control treatment to those produced during Cr-Wr. The analysis showed a reduction from 33% to 27% in sRNAs with a size of 20 nt in wheat during Cr-Wr compared to control wheat roots, while no difference was found between Δ*dcl1*-Wr or Δ*dcl2*-Wr and Cr-Wr (**Figure 8A**). The correlation between mRNA mapping and antisense sRNA mapping was analyzed on every transcript to identify their putative gene targets. On average, the number of transcriptome reads mapping to a transcript remained the same independent of the number of antisense sRNAs mapping to the same transcript (**Figure 8C**).

**Figure 8:**
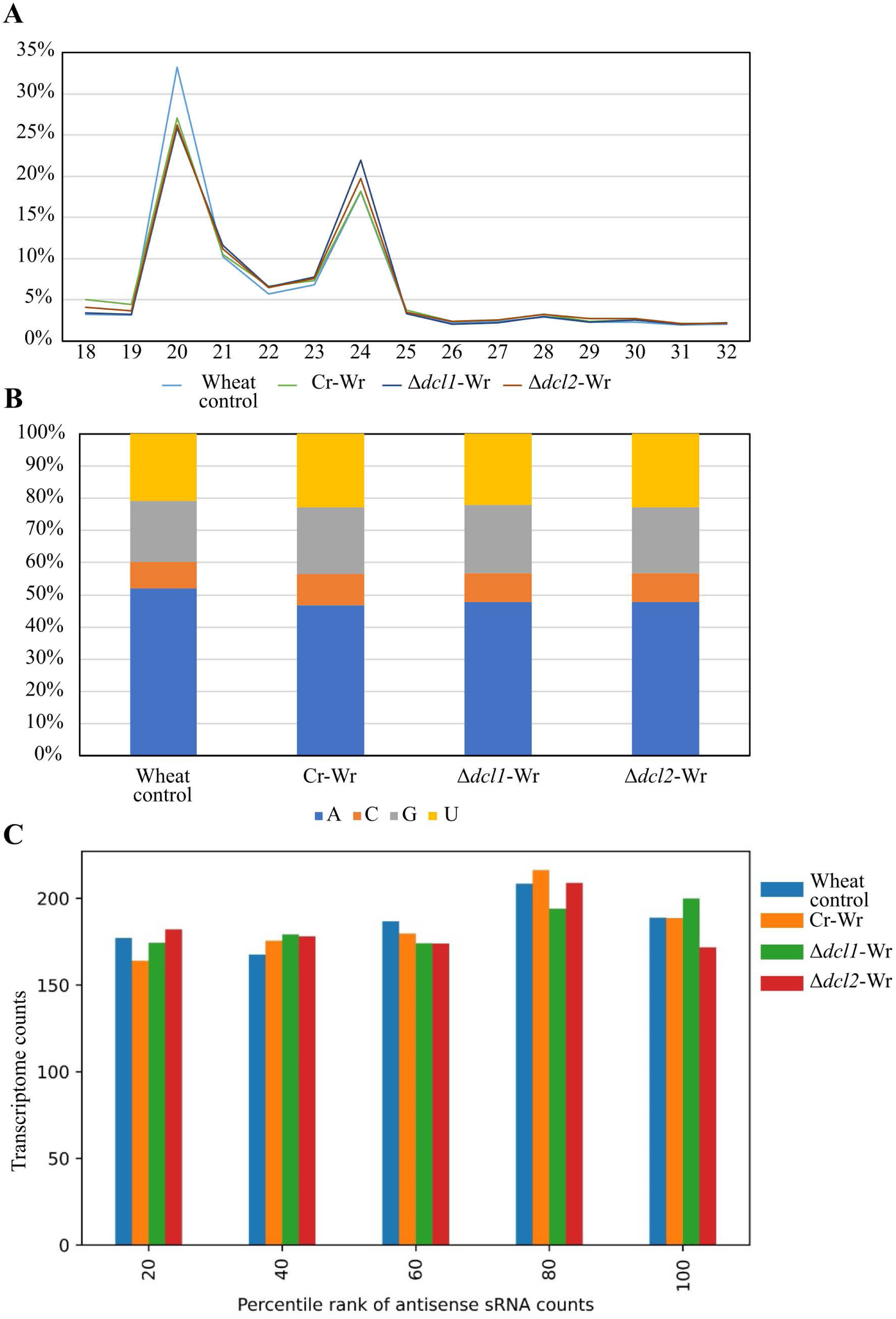
**A:** Read length distribution of sRNA reads mapped to wheat. **B**: 5’ base distribution of sRNA reads mapped to wheat. **C**: Average transcriptome read counts of wheat genes, depending on their percentile rank of antisense sRNA counts. Percentile ranks were assigned to each gene based on its antisense sRNA counts. Genes with an antisense sRNA count of zero were not considered.

We predicted 649 known and seven novel miRNAs in wheat with at least 50 reads (**Supplementary Table 8**), ranging from 19 nt to 23 nt in length. In contrast to the size distribution of total sRNAs, a higher number (47%) of miRNAs were 21 nt in length (**supplementary figure 3A**). Among these, six miRNAs were downregulated during the Cr-Wr compared to wheat control, while three were upregulated. The deletion of *dcl1* in *C. rosea* impacts the miRNA-based response of wheat, with seven miRNAs upregulated and four downregulated during Δ*dcl1*-Wr compared to Cr-Wr interaction. Interestingly, four miRNAs (mir_12061_x13, mir_16010_x2, mir_17532_x1, and mir_19460_x1), which were downregulated during Cr-Wr was found to be upregulated during Δ*dcl1*-Wr (**Table 3).** In the case of Δ*dcl2*-Wr, we detected only three upregulated miRNAs and one downregulated (**Table 3, supplementary figure 3B**).

**Table 3:**
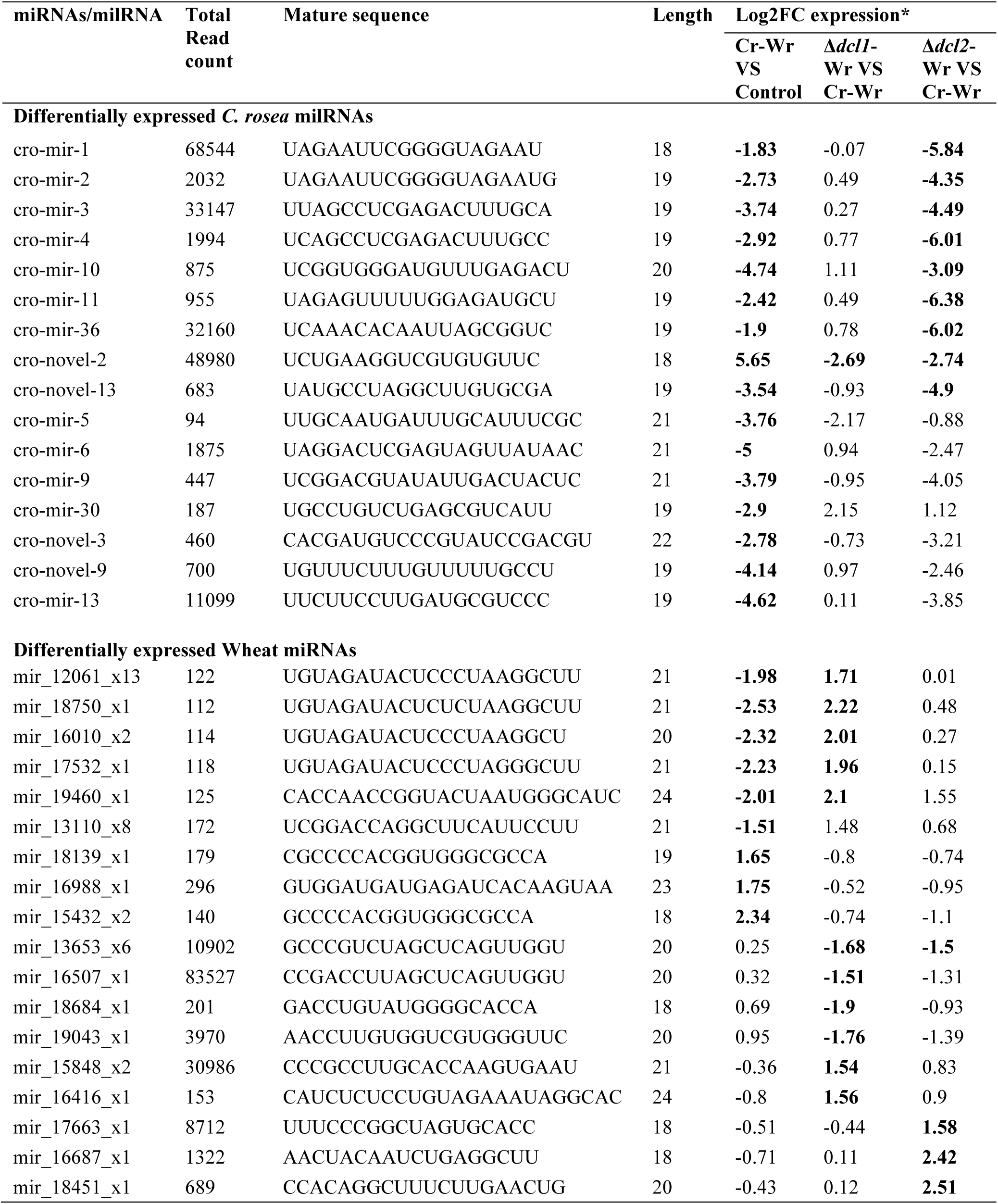
Sequence, length and expression level of differentially expressed milRNAs. Values in bold indicate differential expression with FDR < 0.05.

After target prediction with multiple tools and removal of targets not supported by opposite expression (that is, to consider a transcript as putatively targeted by a miRNA, it had to be upregulated when the miRNA was downregulated), 24 putative endogenous gene targets were identified for seven differentially expressed wheat miRNAs (**supplementary figure 3C; Supplementary Table 9**). However, only four of the targets showed a significant Spearman correlation (Spearman correlation ≤ - 0.77) lesser than -0.4 between target mRNA and targeting miRNA counts (**Table 4**).

**Table 4:** Putative milRNAs targeting endogenous transcripts of interest. All putative targets have been predicted by at least two different target prediction tools, and they show opposite expressions to the targeting milRNAs (a target needs to be upregulated when the targeting milRNA is downregulated). Log2Fc values are in bold when significant (adjusted p-value < 0.05).

| miRNAs/milRNA | Gene target | Spearman correlation | Transcript Expression Log2Fc |  |  | Target gene family | Function |
| --- | --- | --- | --- | --- | --- | --- | --- |
|  |  |  | Cr-Wr | *Δdcl1-Wr* | *Δdcl2-Wr* |  |  |
| Endogenous targets, *C. rosea* |  |  |  |  |  |  |  |
| cro-mir-5 | CRV2T00000691_1 | -0.78 | 0.49 | 0.70 | 0.44 | Transcription factor | Gene expression regulation |
| cro-mir-36 | CRV2T00000889_1 | -0.83 | 2.03 | 1.74 | 1.98 | Transcription factor | Cell integrity, bud morphology and polarization of the actin cytoskeleton |
| cro-mir-3 | CRV2T00002880_1 | -0.69 | 2.71 | 2.51 | 2.68 | Transcription factor | Gene expression regulation |
| cro-mir-10 |  | -0.71 |  |  |  |  |  |
| cro-novel-3 | CRV2T00015685_1 | -0.62 | 0.89 | 0.50 | 0.78 | Transcription factor | Gene expression regulation |
| cro-mir-2 | CRV2T00015018_1 | -0.58 | 1.26 | 0.65 | 0.61 | NRPS | Secondary metabolism |
| cro-mir-30 | CRV2T00004939_1 | -0.77 | 2.71 | 1.98 | 1.78 | MFS transporter | Multidrug resistance |
| cro-mir-10 | CRV2T00006296_1 | -0.74 | 0.31 | -0.07 | 0.39 | ABC transporter | Multidrug resistance |
| cro-mir-11 | CRV2T00006038_1 | -0.59 | 0.44 | 0.50 | 0.47 | Serine/threonine-protein phosphatase PP1 | Cell integrity, bud morphology and polarization of the actin cytoskeleton |
| cro-mir-36 | CRV2T00004592_1 | -0.63 | 8.29 | 7.43 | 8.03 | Pectinesterase, CAZyme CE8+PL1+CBM13 | Plant cell wall degradation |
| cro-novel-9 | CRV2T00007437_1 | -0.59 | 6.29 | 6.07 | 6.35 | Acetyl-coenzyme A synthetase | Acetyl-coenzyme A synthesis |
| cro-novel-9 | CRV2T00013751_1 | -0.61 | 10.05 | 9.52 | 9.62 | Uncharacterized | Unknown |
| cro-mir-36 | CRV2T00015306_1 | -0.81 | 8.88 | 8.37 | 9.18 | CAZyme PL3, probable apoplastic effector | Plant cell wall degradation |
| cro-mir-5 | CRV2T00017422_1 | -0.8 | 9.73 | 9.10 | 8.22 | Probable apoplastic effector | None |
| cro-mir-13 | CRV2T00017982_1 | -0.58 | 6.55 | 4.95 | 5.35 | MFS transporter | Tartrate transporter |
| cro-mir-5 | CRV2T00018464_1 | -0.6 | **6.41** | **6.58** | **6.96** | CAZyme PL3_2, probable apoplastic effector | Plant cell wall degradation |
| cro-mir-5 | CRV2T00018880_1 | -0.59 | **7.71** | **7.46** | **6.12** | CAZyme AA9, probable apoplastic effector | Plant cell wall degradation |
| cro-mir-13 | CRV2T00017601_1 | -0.7 | 0.68 | **2.72** | **4.06** | HlyIII-domain-containing protein | Unknown |
| **Endogenous targets, wheat** |  |  |  |  |  |  |  |
| mir_15432_x2 | TraesCS1D01G107100.1 | -0.77 | **-1.17** | -0.52 | -0.20 | Mixed-linked glucan synthase 8 | Cellulose biosynthesis |
| mir_15848_x2 | TraesCS3D01G258300.1 | -0.83 | 0.32 | -0.56 | -0.37 | ABC transporter | Unknown |
| mir_16507_x1 | TraesCS4D01G102700.1 | -0.82 | -0.21 | 0.38 | 0.42 | L-type lectin receptor kinases | Sensing pathogens and activating defense responses |
| mir_15432_x2 | TraesCS2B01G311600.1 | -0.77 | **-0.58** | -0.4 | -0.48 | transport protein Sec24B-like | Vesicle trafficking |
| mir_18139_x1 |  | -0.76 |  |  |  |  |  |
| **Wheat transcripts targeted by *C. rosea* miRNAs** |  |  |  |  |  |  |  |
| cro-mir-4 | TraesCS4A01G455000.1 | -0.61 | 0.18 | **1.03** | **1.27** | Probable glutathione S-transferase GSTU6 | Resistance to heat |
| cro-mir-11 | TraesCSU01G021200.1 | -0.59 | 0.06 | 0.97 | **1.60** | Blue copper protein | Regulation of lignin biosynthesis and jasmonic acid-based defence |
| ***C. rosea* transcripts targeted by wheat miRNAs** |  |  |  |  |  |  |  |
| mir_17532_x1 | CRV2T00016916_1 | -0.82 | **5.63** | **4.00** | **4.51** | PKS | Synthesis of an antimicrobial compound |
| mir_16010_x2 |  | -0.85 |  |  |  |  |  |
| mir_12061_x13 |  | -0.83 |  |  |  |  |  |

Cross-kingdom target prediction identified six potential cross-kingdom gene targets in *C. rosea* for five wheat miRNAs, which showed an inverse relation in the expression between miRNAs and their corresponding gene targets (**Supplementary Table 9).** However, only one gene (CRV2T00016916_1) showed a significant negative correlation (Spearman correlation ≤ - 0.82) and was identified as a gene targets for three wheat miRNAs (**Table 4**). This gene was identified as polyketide synthase gene *pks29,* shown to be involved in the synthesis of antifungal polyketides (Fatema et al. 2018). Our data showed the potential role of sRNA-mediated gene expression regulation in reprograming its genetic machinery *C. rosea* interactions.

### Identification of *C. rosea* miRNAs differently expressed during the interaction with wheat and their potential endogenous and cross-kingdom gene targets

To investigate sRNA-mediated gene expression regulation in *C. rosea* during interactions with wheat roots, sRNA characteristic and expression pattern was analyzed. The analysis of read length distribution showed peaks of *C. rosea* sRNAs with a size of 19 nt (7-10%), 23 nt (5-7%) and 27 nt (10-15%). Moreover, *C. rosea* control had a higher proportion of 30 nt (17%) sRNAs than Cr-Wr. A peak in sRNAs with a size of 20 nt (11%) was recorded in the Δ*dcl1* strains compared to the other situations (**Figure 9A**). The analysis of 5’ terminal nucleotide composition showed a higher proportion of 5’ – end Uracyl (5’ – U) and 5’ end Guanine (5’ – G) in *C. rosea* control, both occupying around 30% of the reads, followed by 5’ Adenine (5’ – A, 25%) and 5’ Cytosine (5’ – C, 15%). However, during the interactions, the 5’ - A proportion increased to 30%, while the 5’ – U decreased to 25%. The 5’ base distribution was also affected by the deletion of the *dcl* genes, with a reduced proportion of sRNA reads with 5’ – U (20%) and an increased proportion of 5’ C (20%) during Δ*dcl*-Wr (**Figure 9B**). The gene targets of these sRNAs were predicted by mapping to the transcripts. The average number of antisense sRNA reads mapped to a *C. rosea* transcript did not correspond on average with a reduced expression (**Figure 9C**).

**Figure 9:**
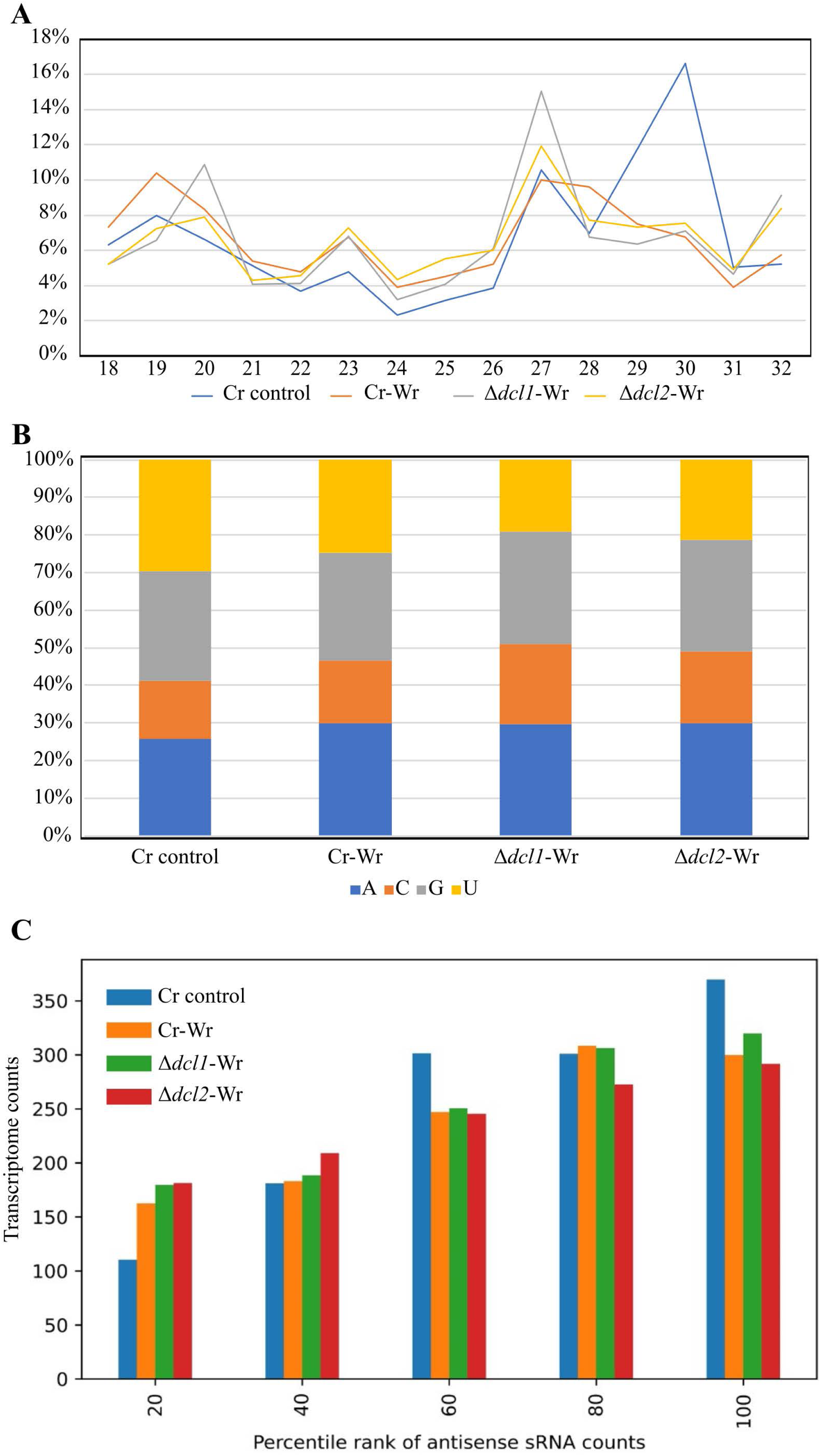
**A**: Read length distribution of sRNA reads mapped to wheat. **B**: 5’ base distribution of sRNA reads mapped to *C. rosea*. **C**: Average transcriptome read counts of *C. rosea* genes, depending on their percentile rank of antisense sRNA counts. Percentile ranks were assigned to each gene based on its antisense sRNA counts. Genes with an antisense sRNA count of zero were not considered.

Our analysis identified 16 known and five novel milRNAs (**Supplementary Table 8**), almost half of which were 19 nt in length (**Supplementary Figure 4A**). Of these, 15 milRNAs were downregulated during Cr*-*Wr interaction, compared to *C. rosea* control. Nine milRNAs were downregulated during the Δ*dcl2*-Wr compared to the Cr-Wr, indicating their origin was DCL-dependent, while only one milRNA was downregulated during Δ*dcl1*-Wr (**Table 3, Supplementary Figure 4B**). Differentially expressed *C. rosea*’s milRNAs had 480 putative endogenous targets (**supplementary table 9**). 320 showed a significant inverse correlation between gene target expression and their corresponding milRNAs (Spearman correlation > - 0.4).

A higher number of endogenous gene targets (405) was predicted for downregulated *C. rosea* milRNAs. These consisted of gene coding for transcription factors (11 genes), 12 putative effectors (12 genes), 19 MFS transporters (19 genes), and two genes each coding for core enzymes of specialized metabolite gene clusters, RNA helicases and chromatin remodelling proteins (**Supplementary Table 9**). Additionally, 11 glycoside hydrolase genes were detected in this dataset, including members of families GH10, GH15, GH26, GH3, GH35, GH43, GH45, GH5 and GH7. On the contrary, only one milRNA (cro-novel-2) was upregulated during Cr-Wr, compared to the control, and it was predicted to target three genes: a transcription factor, a proteolytic enzyme and a protein involved in guanosine tetraphosphate metabolism (**Supplementary Table 9**).

The nine DCL2-dependent milRNAs were predicted to have 81 gene targets, including three genes coding for transcription factors (CRV2T00000423_1, CRV2T00004800_1, CRV2T00015277_1), three effectors (CRV2T00014512_1, CRV2T00019066_1, CRV2T00019646_1), three MFS transporters (CRV2T00008216_1, CRV2T00013299_1, CRV2T00013418_1) and one protein (CRV2T00007349_1) involved in histone acetylation.

We identified 38 putative gene targets in wheat (cross-kingdom RNA silencing) for eight milRNAs of *C. rosea*. Of these transcripts, 29 were putatively targeted by seven DCL2-dependent *C. rosea* milRNAs. Among these targets, genes coding for a glutathione transferase GSTU6, a blue copper protein, a GDSL esterase/lipase and a LRR receptor-like serine/threonine-protein kinase are the most significantly upregulated gene targets. The other gene targets consisted of four xyloglucan endotransglycosylases, three peroxidases 11, one pathogen-related protein, a probable LRR receptor-like serine/threonine-protein kinase, one transcription factor and six histone-like proteins (**Table 4; Supplementary Table 9**).

## Discussion

Plant-beneficial fungi, for example those belonging to the genus *Trichoderma* are shown to trigger plant defense response during interaction with plant host (Coppola et al., 2019; De Palma et al., 2019). However, the interplay between the biocontrol fungus *C. rosea* and its host roots remains elusive. In this study, we investigated the interactions between the fungal biocontrol agent *C. rosea* and wheat roots, focusing on the transcriptional changes and their regulation that occur during these interactions. We also explored the role of DCLs in regulating gene expression in *C. rosea* during these interactions and its cross-kingdom effect. Previous gene expression studies using real-time qPCR have indicated the ability of *C. rosea* to induce the defence response of plant hosts (Roberti et al. 2008; Kamou et al. 2020; Dubey et al. 2020; Sun et al. 2020). In the present work, we showed that wheat reacted to the interaction with the fungus by upregulating many stress resistance genes and downregulating genes involved in cell wall expansion and biosynthesis. We hypothesize that the transcriptional response of wheat is at least partly caused by a *C. rosea* plant cell wall degrading activity. This is supported by the gene enrichment analysis showing *C. rosea* upregulated genes Cr-Wr were enriched in terms related to polysaccharide catabolism, cell wall modification, localization, transmembrane transport, and extracellular region. This suggests that C. *rosea*-produced enzymes are able to degrade the plant cell wall and transports them to the fungal cell. As a reaction to *C. rosea-* mediated degradation, wheat reprogramed its genetic machinery for the increased defense-response and decreased cell growth by upregulating stress-related genes and, at the same time, downregulating gens associated with development. This growth-defense trade-off is a well-known phenomenon for resource allocation in plants to optimize fitness during host-microbe interactions and stress (Huot et al., 2014).

Many *Trichoderma* spp. share a similar ecological niche to *C. rosea* and are similarly studied for their biocontrol capabilities. Despite this, we observed that the effect of *C. rosea* on wheat roots is quite different to what was observed with *Trichoderma harzianum* by Rubio et al. (2019). The ethylene pathway was induced by both biocontrol agents, with the upregulation of ethylene-responsive transcription factor RAP2-3 like TraesCS1A01G231200.1, and chymotrypsin inhibitors also upregulated during the interaction with both organisms (Rubio et al. 2019). Wound-responsive gene CS2B01G561300.1 was also upregulated in both situations, but *T. harizanum* was also able to activate genes linked to the abscisic acid response (Rubio et al. 2019), while *C. rosea* induced the upregulation of genes encoding for LEA proteins, dehydrins, vicilin-like storage proteins and lectins, and several known disease-resistance proteins. Another difference was that *T. harzianum* induced the downregulation of very few genes (25% of the upregulated ones), while in the current study, the number of wheat genes upregulated and downregulated in Cr-Wr was similar. A Gene coding for Expansin (TraesCS4A01G034300.1) was downregulated in both experiments (Rubio et al. 2019), but nine other proteins of the same class were also affected during *C. rosea* interaction with wheat. Unfortunately, there are no studies on the *Trichoderma* transcriptomics response to wheat roots. Still, Morán-Diez et al. (2015) studied the response of *Trichoderma virens* to maize roots, observing how the fungus upregulated many transporter genes as well as glycoside hydrolases (Morán-Diez et al. 2015). After prediction with dbCAN2 (Zhang et al. 2018), however, only one of the *T. virens* genes upregulated during the response to maize roots (gene 11696) belong to the AA9 CAZyme class, which was the one with more upregulated member in *C. rosea*. This is not surprising, as *Clonostachys* spp. are known to have a much higher number of AA9 in their secretome, compared with *Trichoderma* spp. (Piombo et al. 2023). In summary, the result highlighted that the interaction mechanisms between beneficial fungi and host plants depend on the interactions organisms and the experimental conditions.

The interaction between wheat and *C. rosea* is heavily affected by deleting *dcl* genes in the latter. We identified 65 wheat stress-responsive genes that are upregulated in Cr-Wr but not in either Δ*dcl1*-Wr or Δ*dcl2*-Wr, suggesting an essential role of DCL-dependent regulation in the capacity of the fungus to induce defense reactions in the plant **(Figure 6**). Such defense reaction could be caused by *C. rosea* cell wall degrading enzymes, many of which are upregulated in Cr-Wr but not in Δ*dcl1*-Wr or Δ*dcl2*-Wr. Furthermore, the interaction with the WT fungus, but not with the mutants, induced in wheat a downregulation of genes having a role in the loosening of the plant cell wall, in the form of expansin-like proteins and a peroxidase-57, whose overexpression increases cuticle permeability in *Arabidopsis thaliana* (Survila et al. 2016). The mutants will, therefore, interact with a looser and more permeable plant cell wall while simultaneously needing to cope with less plant defense reactions. This could explain the increased root colonization capacity of the Δ*dcl2* mutant compared to *C. rosea* WT.

The response of wheat and *C. rosea* to each other appears to be partly milRNA-dependent. Six wheat miRNAs and 15 *C. rosea* milRNAs were downregulated during plant-fungus interaction compared to the respective control conditions. In comparison, only three plant miRNAs and one fungal milRNAs were upregulated when the two organisms were interacting. This suggests that many milRNAs necessary to regulate the interaction are constitutively expressed in both organisms but downregulated during the interaction, enabling the expression of transcripts that they normally negatively regulate.

The fungal milRNAs downregulated during the interactions were predicted to target five transcription factors. One core NRPS of a specialised metabolite gene cluster was also targeted (CRV2T00015018_1). Other targets of interest were two transmembrane transporters, 1 of which (CRV2T00004939_1) was overexpressed in *C. rosea* during interaction with the plant pathogens *F. graminearum* or *B. cinerea* (Nygren et al. 2018), as well as the serine/threonine-protein phosphatase PP1 (CRV2T00006038_1), involved in cell integrity, bud morphology and polarization of the actin cytoskeleton (Andrews and Stark 2000). The most upregulated eight targets (log2FC > 5.75), however, were a putative apoplastic effector, an AA9 CAZyme, a PL1 pectinesterase, two pectate lyases of the PL3 family, an Acetyl-coenzyme A synthetase, an MFS transporter and one uncharacterized protein. CAZyme class AA9 is known to increase the action of other cellulases (Harris et al. 2010). The PL1 pectinesterase and 2 PL3 enzymes are also involved in the degradation of plant tissues, and heterologous expression of PL1 and PL3 enzymes is known to confer resistance to *Erwinia carotovara* in potatoes (Wegener et al. 1996; Wegener 2002; Wegener and Olsen 2004). These proteins, negatively regulated by milRNAs in the control condition but not during plant-fungus interaction, could participate in the defense induction on wheat by partially degrading the plant cell wall **(Table 4**).

Genes of interest were also targeted by milRNAs upregulated during the plant-fungus interaction. In particular, wheat mir_15432_x2 was predicted to target a mixed-linked glucan synthase eight involved in cellulose synthesis (TraesCS1D01G107100.1), suggesting wheat uses RNA silencing to reduce cellulose production during the interaction with *C. rosea*. This miRNA was also identified as upregulated in the wheat cultivar Zhengyin 1 after dehydration stress (Ma et al. 2015). On the fungal side, however, only the milRNA cro-novel-2 was more expressed in the plant-fungal interaction than in control. It had only three putative targets, none showing significant anti-correlation with the targeting milRNA.

The silencing of *dcl* genes in *C. rosea* altered the effect of the fungus on wheat, and the plant reacted to the mutants by producing different miRNAs concerning how it responded to the WT. Plant miRNA mir_15848_x2 was upregulated in the response to the Δ*dcl1* mutant, and it was predicted to target the ABC transporter TraesCS3D01G258300.1. In contrast, three different miRNAs (mir_17532_x1, mir_16010_x2 and mir_12061_x13) were all predicted to target the *C. rosea* transcript CRV2T00016916_1, downregulated during the interaction between the mutant and the plant in comparison to the WT-wheat interaction. This transcript encodes a polyketide synthase known as PKS29, producing an unknown compound with a proven antimicrobial activity (Fatema et al. 2018). As already seen in a previous study (Piombo et al. 2021), the deletion of *dcl2* affected a higher number of *C. rosea* milRNAs (nine) than that of *dcl1*. Moreover, the only milRNA downregulated in the Δ*dcl1*-wheat interaction, compared to the WT-wheat interaction, was also downregulated in the Δ*dcl2* mutant.

DCL2-dependent fungal milRNAs were also predicted to target 28 plant genes. The ones showing a more evident opposite expression to the targeting milRNAs are a blue copper protein and glutathione transferase GSTU6. A blue copper protein affects the regulation of lignin biosynthesis and jasmonic acid-based defence in cotton (Zhu et al. 2018), while Glutathione S-transferase GSTU6 has a unique SNP in heat-tolerant rice varieties and is less expressed in these varieties than in heat sensitive ones (Zhou et al. 2019), suggesting that downregulation of the gene, such as the one observed in Cr-Wr but not in Δ*dcl2*-Wr mutant, could increase heat resistance in wheat. All of this suggests that *C. rosea* could use milRNAs to affect the expression of wheat genes involved in resistance to both abiotic and biotic stresses.

## Conclusions

In conclusion, the interaction between wheat and the biocontrol fungus *C. rosea* is a complex and dynamic process that involves the expression of numerous genes and the regulation of miRNAs in both the plant and the fungus. This interaction triggers a cascade of molecular events in wheat, leading to the activation of stress resistance genes and the modulation of cell wall-related processes, suggesting a trade-off between defense and growth. *Clonostachys rosea*, in turn, showed altered expression of genes involved in carbohydrate catabolism, membrane transport, and the production of effector molecules. In addition, the study explores the effects of DCL (Dicer-like) gene deletions in *C. rosea* and their impact on root colonization, as well as the transcriptomic responses of both *C. rosea* and wheat during their interactions. Notably, deleting DCL genes in *C. rosea* alters its impact on wheat, resulting in differential gene expression profiles of genes associated with the growth and resistance to abiotic stresses. Deletion of *dcl2* had a more pronounced effect on wheat gene expression compared to *dcl1* deletion, which is in line of the increased root colonization ability of DCL2 deletion strains. The study identified candidate miRNAs in what and milRNAs in *C. rosea* and their potential endogenous and cross-kingdom gene targets. These were differentially expressed during their interactions, suggesting a complex network of sRNA-mediated gene regulation during the interactions.

This study underscores the importance of sRNAs as crucial players in the intricate molecular interplay between wheat and *C. rosea,* regulating gene expression potentially at both endogenous and cross-kingdom levels. Further exploration of these sRNA-mediated mechanisms and functional characterization of candidate miRNAs and milRNAs can provide valuable insights into the biocontrol potential of *C. rosea*. It may help to develop novel strategies for enhancing plant resistance to both biotic and abiotic stressors in agriculture. Further research into the specific roles of identified genes and miRNAs/milRNAs in this interaction will be essential for a deeper understanding of the underlying mechanisms and their potential applications in crop protection and stress management.

## Materials and methods

### Sample preparation for RNA sequencing

Surface sterilized wheat seeds were germinated on sterilized moist filter paper placed in nine-cm petri plates (five seeds per plate) following the procedures described before (Dubey et al. 2020). Surface sterilized germinated as described in the gene expression section. Three days old wheat seedlings were inoculated by dipping the roots for three minutes in *C. rosea* spore suspensions (1e+07 spore/ml) in sterile condition, transferred back to the filter paper in petri plates and incubated at 20°C as described before (Dubey et al., 2021). Roots were harvested seven dpi and snap-frozen in liquid nitrogen. Three biological replicates with five seedlings per replicate were used for each treatment.

### Root colonization assay

An experimental setup was done as described above to quantify the root colonisation. Root colonization was determined five dpi by quantifying the DNA level of *C*. *rosea* strains in wheat roots using qPCR (Dubey et al. 2013). The actin gene was used as the target gene for *C*. *rosea*, and *Hor1* (Nicolaisen et al., 2009) was used as the target gene for wheat (Dubey et al., 2021). Root colonization was expressed as the ratio between *actin* and *Hor1*.

### RNA extraction library preparation and sequencing

Total RNA was extracted using the mirVana miRNA isolation kit following the manufacturer’s protocol (Invitrogen, Waltham, MA). The RNA quality was analyzed using a 2100 Bioanalyzer Instrument (Agilent Technologies, Santa Clara, CA), and concentration was measured using a Qubit fluorometer (Life Technologies, Carlsbad, CA). For sRNA and mRNA sequencing, the library was prepared and paired-end sequenced at the National Genomics Infrastructure (NGI) in Stockholm, Sweden. The sRNA library was generated using the TruSeq small RNA kit (Illumina, San Diego, CA), while the mRNA library was generated using the TruSeq Stranded mRNA poly-A selection kit (Illumina, San Diego, CA). The sRNA and mRNA libraries were sequenced on one NovaSeq SP flowcell with a 2×50 bp reads and NovaSeqXp workflow in S4 mode flow cell with 2×151 setup, respectively, using Illumina NovaSeq6000 equipment at NGI Stockholm. The Bcl to FastQ conversion was performed using bcl2fastq_v2.19.1.403 from the CASAVA software suite (Hosseini et al. 2010). The quality scale used was Sanger / phred33 / Illumina 1.8+.

### Mapping and differential expression analyses

Adapter and quality trimming was done for sRNA and mRNA reads using bbduk v. 38.9 (Bushnell 2019), and quality was then checked using fastqc (Andrews 2010). The options used for bbduk were: ktrim=r k=23 mink=11 hdist=1 tpe tbo qtrim=r trimq=10 For mRNAs, reads were mapped to the *C. rosea* IK726 genome (GCA_902827195) and the IWGSC ‘Chinese Spring’ genome assembly (Consortium 2014) using the splice-aware aligner STAR (Dobin et al. 2013) with default parameters and the option “–outFilterMultimapNmax 40”. Reads mapping to both genomes were then excluded with an ad hoc pipeline using samtools v. 1.9 (Danecek et al. 2021) and Picard tools v. 2.18.29 (http://broadinstitute.github.io/picard/). The number of reads mapping to each gene was then evaluated using featureCounts v. 2.0.1 (Liao et al. 2014), and differential expression was determined with the DESeq2 R package v. 1.28.1 (Love et al. 2014) using a minimal threshold of 1.5 for log2(FC) and 0.05 for FDR adjusted p-value.

Normalized expression values were obtained from DESeq2 and used to perform a coexpression analysis with WCGNA (Langfelder and Horvath, 2008) using only differentially expressed genes. The soft-thresholding power was 6 for *C. rosea* genes and 16 for wheat genes, and the function “binarizeCategoricalVariable” was used to convert the categorical variables into numerical ones.

AgriGO v. 2 (Tian et al. 2017) was used to determine enriched GO terms in the differentially expressed genes, using a Fisher test with Yekutieli (FDR under dependency) adjustment method. The adjusted p-value threshold was set at 0.05, and enriched biological processes were visualized with REVIGO (Supek et al. 2011).

Reformat.sh v. 38.9 (Bushnell 2019) was used only to retain sRNA reads between 18 bp and 32 bp in length, and rRNAs, tRNAs, snRNAs and snoRNAs were removed from the dataset using SortMeRNA v. 4.2.0 (Kopylova et al. 2012) using as references RNA sequences downloaded from SILVA and the NRDR database (Paschoal et al. 2012; Quast et al. 2013). The filtered sRNA reads were then mapped to the *C. rosea* and wheat transcriptomes using bowtie v. 0.12.9 (Langmead 2010) using options:

-S -k 101 -n 2 -l 18 -m 200 –best –strata

In the case of wheat, only High Confidence transcripts (https://urgi.versailles.inra.fr/download/iwgsc/IWGSC_RefSeq_Annotations/v1.0/) were used, and the options “-n 3” were added to compensate for the fact that the sequenced genome was of a different cultivar from the one used for the experiment.

FeatureCounts v. 2.0.1 was used to quantify the reads mapping to each transcript, only counting antisense mappings, and normalized sRNA-mapping values were obtained for each gene using DESeq2 (Love et al. 2014). Spearman and Pearson correlations were then calculated between the sRNA mapping and mRNA mapping of each differentially expressed gene, and the anticorrelation was classified as “moderate” when the Spearman correlation was between -0.4 and -0.7, “strong” when it was between -0.7 and -0.9 and “very strong” if it was less than -0.9.

### Functional annotation

The function of *C. rosea* genes was determined according to functional annotation performed in previous publications (Piombo et al. 2021), while differentially expressed wheat genes had their domains predicted with InterProScan (Jones et al. 2014), and they were compared with the NCBI database through BLAST (Altschul et al. 1990). Additionally, effectorP was used to predict *C. rosea* effector-like proteins (Sperschneider and Dodds 2021).

### milRNA prediction, differential expression and target prediction

Known milRNA sequences of *C. rosea* were retrieved from previous publications (Piombo et al. 2021), while known wheat miRNAs were retrieved from the Wheat miRNA web portal and 16iRbase (Remita et al. 2016; Kozomara et al. 2019). Novel milRNAs of *C. rosea* were predicted with MiRDeep2 v. 2.0.1.3 using default parameters (Friedländer et al. 2012; Kuang et al. 2019), while novel wheat miRNAs were predicted using MiRDeep-P2 v. 1.1.4.

The presence of each milRNA/miRNAs was quantified in each sample by counting their occurrence in the clean reads file allowing for one mismatch using agrep (Ahmad et al. 2006). Target prediction was done using the plant-based tools psRNATarget, Targetfinder, psROBOT and TAPIR, using the latter two through the sRNA toolbox (Bo and Wang 2005; Bonnet et al. 2010; Wu et al. 2012; Rueda et al. 2015; Dai et al. 2018) and milRNA-target couples predicted by at least two tools were retained. Self-targets of *C. rosea* milRNAs were also predicted with the animal-based tools PITA, Miranda, TargetSpy and simple seed analysis, all used through the sRNA toolbox, and milRNA-target couples predicted by at least three tools were retained (Enright et al. 2003; Kertesz et al. 2007; Sturm et al. 2010; Rueda et al. 2015). Afterwards, we only retained milRNA-target couples showing opposite expression between the milRNA and the putative target (if one is up-regulated in a specific condition, then the other needs to be down-regulated). For this filtering step, we used DESeq2 v. 1.28.1 (Love et al. 2014) with a minimal threshold of 1.5 for log2(FC) and 0.05 for FDR-adjusted p-value for milRNAs, while for putative targets, we set a threshold of 0.05 for FDR adjusted p-value but no threshold for log2(FC). Additionally, for each target, we calculated the Spearman and Pearson correlation between the miRNA counts and the target mRNA counts, and we compared it to the average correlation of the miRNA with any other transcript of the same organism. As done in a previous study (Wang and Li 2009), we used the Wilcoxon rank sum test with a p-value threshold of 0.1 to determine if the anti-correlation between milRNA and target was significantly higher than the one between the milRNA and the average transcript. Only targets with significant Spearman anticorrelation were examined to determine targets of interest, but the values for both Spearman and Pearson anticorrelation are nevertheless available in **Supplementary Table 8**.

## Acknowledgements

This work was financially supported by the Department of Forest Mycology and Plant Pathology; Swedish Research Council for Environment, Agricultural Sciences and Spatial Planning (FORMAS; grant number 2018-01420; 2021-01461), and Carl Tryggers Stiftelse för Vetenskaplig Forskning (CTS 19: 82). RV is supported by FORMAS (2019-01316), Carl Tryggers Stiftelse för Vetenskaplig Forskning (CTS 20: 464) and The Crafoord foundation (20200818). The authors acknowledge support from the National Genomics Infrastructure in Stockholm funded by Science for Life Laboratory, the Knut and Alice Wallenberg Foundation and the Swedish Research Council, and SNIC/Uppsala Multidisciplinary Center for Advanced Computational Science for assistance with massively parallel sequencing and access to the UPPMAX computational infrastructure.

## Supplementary information

**Supplementary Table 1:** Sample-by-sample mRNA and sRNA mapping results on wheat and *C. rosea*.

**Supplementary Table 2:** Transcripts predicted to be differentially expressed in this study. Analysis carried out through DESeq2 v. 1.28.1 with default parameters. The adjusted p-value threshold was fixed at 0.05, and minimum log2(FC) was set at 1.5.

**Supplementary Table 3:** Gene ontology terms (GOs) enriched in wheat and *C. rosea* genes upregulated or downregulated during Cr-Wr. The analysis was carried out with AgriGO v. 2 using a Fisher test with Yekutieli (FDR under dependency) adjustment method and adjusted p-value threshold set at 0.05.

**Supplementary Table 4:** Wheat genes of interest upregulated or downregulated during the interaction with *C. rosea* WT.

**Supplementary Table 5:** Top 20 highly upregulated or downregulated wheat genes during *C. rosea*- wheat interactions compared to wheat control.

**Supplementary Table 6:** Top 20 highly upregulated wheat genes or downregulated *C. rosea* genes during the interactions with wheat roots.

**Supplementary Table 7:** Differentially expressed wheat genes with a role in cell wall synthesis or modification, resistance, or induction of defense reactions, as well as *C. rosea* CAZymes or effectors. All the genes were upregulated or downregulated in Cr-Wr but not in Δ*dcl1*-Wr and Δ*dcl2*-Wr. Log2Fc values are in bold when significant (adjusted p-value < 0.05).

**Supplementary Table 8:** Sequence, length, and expression level of detected expressed milRNAs. Note that in this analysis, the conditions “Δ*dcl1*-Wr” and “Δ*dcl2*-Wr” were compared with “Cr-Wr” and not to the control in the differential expression analysis. This way, the conditions involving mutants (Δ*dcl1*-Wr and Δ*dcl2*-Wr) were compared directly with the same condition with the WT (Cr-Wr) rather than with *C. rosea in vitro* or non-inoculated wheat.

**Supplementary Table 9:** Transcripts predicted to be targeted by putative milRNAs. All putative targets have been predicted by at least two target prediction tools, and they show opposite expressions to the targeting milRNAs (a target needs to be upregulated when the targeting milRNA is downregulated). Note that in this analysis, the conditions “Δ*dcl1*-Wr” and “Δ*dcl2*-Wr” were compared with “Cr-Wr” and not to the control in the differential expression analysis. This way, the conditions involving mutants (Δ*dcl1*-Wr and Δ*dcl2*-Wr) were compared directly with the same condition with the WT (Cr-Wr) rather than with *C. rosea in vitro* or non-inoculated wheat.

## Notes

### Competing Interest Statement

The authors have declared no competing interest.

