## Supplemental Table 5 for "Dicer-mediated RNA silencing is the key regulatory mechanism in the biocontrol fungus *Clonostachys rosea*-wheat interactions"

Table S5: Top 20 highly upregulated or downregulated wheat genes during *Clonostachys rosea* -wheat interactions (Cr-Wr) compared to wheat control.

| Transcript | Log2Fc | Target gene | Characterized/putative function |
| --- | --- | --- | --- |
| **Top 20 highly upregulated wheat genes during Cr-Wr** | | | |
| Traescs5a01g329100.1 | 8.5 | Defensin-like-1 | Antifungal activity |
| Traescs4d01g262800.1 | 7.0 | Vicilin-like seed storage protein | Inhibiting spore germination and growth of plant pathogens |
| Traescs1b01g101400.1 | 6.7 | Ubiquitinyl hydrolase 1 | Deubiquitination |
| Traescsu01g122200.1 | 6.1 | Dehydrin | Response to dehydration and cold |
| Traescs4d01g056000.1 | 5.7 | 7-methyl-GTP pyrophosphatase-like | None |
| Traescs4b01g106100.1 | 5.4 | Fusarium resistance orphan protein | Resistance to *F. graminearum* |
| Traescs7a01g416600.1 | 5.3 | Disease-resistance protein RGA5-like | Resistance to pathogens |
| Traescs4a01g244800.1 | 5.0 | Tetratricopeptide repeat-like protein | Immune response |
| Traescs1a01g000200.1 | 5.0 | Latent-transforming growth factor beta-binding protein 3 | None |
| Traescs4a01g250900.1 | 5.0 | Dehydrin | Response to dehydration and cold |
| Traescs5b01g180700.1 | 5.0 | F-box protein-like protein | Resistance to biotic and abiotic stress |
| Traescs2b01g132500.1 | 4.9 | UDP-glucosyltransferase | Abiotic stress tolerance |
| Traescs1a01g001300.1 | 4.8 | Protein TAR1-like | Auxin biosynthesis |
| Traescs6b01g117800.1 | 4.8 | Uncharacterized | None |
| Traescs4b01g262700.1 | 4.8 | Vicilin-like seed storage protein | Inhibiting spore germination and growth of plant pathogens |
| Traescs7a01g269000.1 | 4.7 | Uncharacterized | None |
| Traescs7d01g119300.1 | 4.7 | Uncharacterized | None |
| Traescs1a01g001400.1 | 4.7 | Uncharacterized | None |
| Traescs1b01g248400.1 | 4.6 | Threonine--tRNA ligase | Protein translation |
| Traescs6d01g260500.1 | 4.6 | Protein NRT1/ PTR FAMILY 5.1 | Oligopeptide transport |
| **Top 20 highly downregulated wheat genes during Cr-Wr** | | | |
| Traescs6b01g246400.1 | -6.5 | RNA-binding protein | RNA-binding protein |
| Traescs5d01g446900.1 | -4.7 | Pathogenesis-related protein 1 | Resistance to pathogens |
| Traescs5d01g488700.1 | -2.8 | Chalcone synthase | Resistance to biotic and abiotic stress |
| Traescs5d01g488800.1 | -2.8 | O-methyltransferase | None |
| Traescs5d01g488600.1 | -2.8 | Chalcone synthase | Resistance to biotic and abiotic stress |
| Traescs5d01g286000.1 | -2.7 | Auxin-induced in root cultures protein 12 | None |
| Traescs3b01g152600.1 | -2.7 | Glucuronosyltransferase | None |
| Traescs2d01g498700.1 | -2.6 | Probable xyloglucan endotransglucosylase/hydrolase | Expansive growth of plant cell walls |
| Traescs2d01g498600.1 | -2.6 | Xyloglucan endotransglucosylase/hydrolase | Expansive growth of plant cell walls |
| Traescs4b01g052100.1 | -2.6 | Probable protein kinase | None |
| Traescs2d01g026800.1 | -2.6 | Serine/threonine-protein phosphatase 7 | None |
| Traescs3a01g084100.1 | -2.6 | Transmembrane protein 45B-like | None |
| Traescs4a01g463400.1 | -2.6 | O-methyltransferase | None |
| Traescs1b01g214800.1 | -2.5 | Peroxidase 5-like | Resistance to pathogens |
| Traescs4d01g324300.1 | -2.5 | Uncharacterized | None |
| Traescs2a01g470100.1 | -2.5 | IQ domain-containing protein | None |
| Traescs5d01g488900.1 | -2.4 | O-methyltransferase | None |
| Traescs7a01g566600.1 | -2.4 | Putative NBS-LRR resistance protein | Resistance to pathogens |
| Traescs2a01g429700.1 | -2.4 | Zealexin A1 synthase-like | Phytoalexin synthesis |
| Traescs4b01g309900.1 | -2.4 | Serpin | Inhibition of hypersensitive response |
