## Supplemental Table 6 for "Dicer-mediated RNA silencing is the key regulatory mechanism in the biocontrol fungus *Clonostachys rosea*-wheat interactions"

**Supplementary Table 6:** Top 20 highly upregulated wheat genes or downregulated *C. rosea* genes during the interactions with wheat roots*.*

| **Transcript** | **Log2Fc** | **Target gene family** | **Characterized/putative function** |
| --- | --- | --- | --- |
| **Top 20 *C. rosea* genes highly upregulated during the interactions with wheat roots.** | | | |
| CRV2T00014599_1 | 11.7 | Acetyl xylan esterase CE1+CBM1+CBM2; Effector | Xylan degradation |
| CRV2T00013015_1 | 10.5 | Alpha-L-arabinofuranosidase GH54+CBM13+CBM42 | Hemicellulose degradation |
| CRV2T00005329_1 | 12.5 | CBM1 CAZyme; Effector | None |
| CRV2T00014442_1 | 12.4 | Cellulase GH12; Effector | Cellulose degradation |
| CRV2T00022277_1 | 12.4 | Cellulase GH12; Effector | Cellulose degradation |
| CRV2T00019248_1 | 12.5 | Effector | None |
| CRV2T00009771_1 | 11.3 | Effector | None |
| CRV2T00003184_1 | 10.5 | Effector | None |
| CRV2T00014519_1 | 10.3 | Effector | None |
| CRV2T00010395_1 | 11.3 | Glucuronyl esterase CE15+CBM1 | Degradation of lignocellulose |
| CRV2T00000047_1 | 10.4 | MFS transporter | None |
| CRV2T00012723_1 | 13.6 | Monooxygenase AA9+CBM1; Effector | Initiate breakdown of cellulose |
| CRV2T00013543_1 | 13.4 | Monooxygenase AA9+CBM1; Effector | Initiate breakdown of cellulose |
| CRV2T00008335_1 | 10.2 | Monooxygenase AA9+CBM1; Effector | Initiate breakdown of cellulose |
| CRV2T00003272_1 | 11.4 | Monooxygenase AA9; Effector | None |
| CRV2T00018190_1 | 10.2 | Monooxygenase AA9; Effector | None |
| CRV2T00013064_1 | 10.3 | Oxidoreductase | None |
| CRV2T00003237_1 | 11.9 | Putative aspergillopepsin | Aspartic-type endopeptidase activity |
| CRV2T00009945_1 | 11.8 | Uncharacterized | None |
| CRV2T00014860_1 | 11.5 | Uncharacterized | None |
| **Top 20 *C. rosea* genes most downregulated regulated during the interactions with wheat roots** | | | |
| CRV2T00009702_1 | -9.3 | Uncharacterized | None |
| CRV2T00003716_1 | -9.2 | NRPS | Specialised metabolism |
| CRV2T00009013_1 | -8.4 | Oxidoreductase | None |
| CRV2T00003718_1 | -8.2 | Uncharacterized | None |
| CRV2T00011478_1 | -8 | MFS transporter | Iron transport |
| CRV2T00018816_1 | -7.5 | Effector | None |
| CRV2T00016772_1 | -7.5 | Zinc permease | Zinc transport |
| CRV2T00020557_1 | -7.3 | Transcription factor | None |
| CRV2T00020556_1 | -7.3 | Transcription factor | None |
| CRV2T00017919_1 | -7.3 | Transcription factor | None |
| CRV2T00017382_1 | -7.3 | Oxidoreductase | None |
| CRV2T00003715_1 | -6.9 | MFS transporter | None |
| CRV2T00009227_1 | -6.9 | Superoxide dismutase | Defense against reactive oxygen species |
| CRV2T00009228_1 | -6.8 | Uncharacterized | None |
| CRV2T00003719_1 | -6.7 | Lipase | None |
| CRV2T00015361_1 | -6.7 | Copper transporter | Copper transport |
| CRV2T00011568_1 | -6.5 | Uncharacterized | None |
| CRV2T00013955_1 | -6.4 | Uncharacterized | None |
| CRV2T00019717_1 | -6.4 | Uncharacterized | None |
| CRV2T00011718_1 | -6.4 | Alkaline phosphatase | None |
