## Supplementary figures and images for "Dicer-mediated RNA silencing is the key regulatory mechanism in the biocontrol fungus *Clonostachys rosea*-wheat interactions"

### Supplemental Figure 1

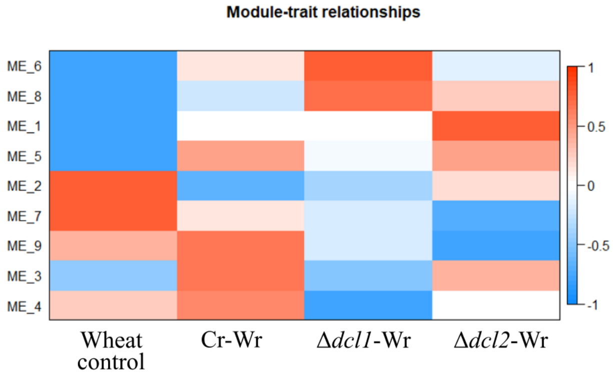

### Supplemental Figure 2

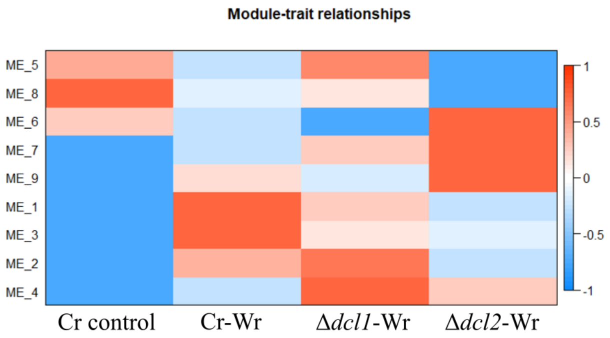

### Supplemental Figure 3

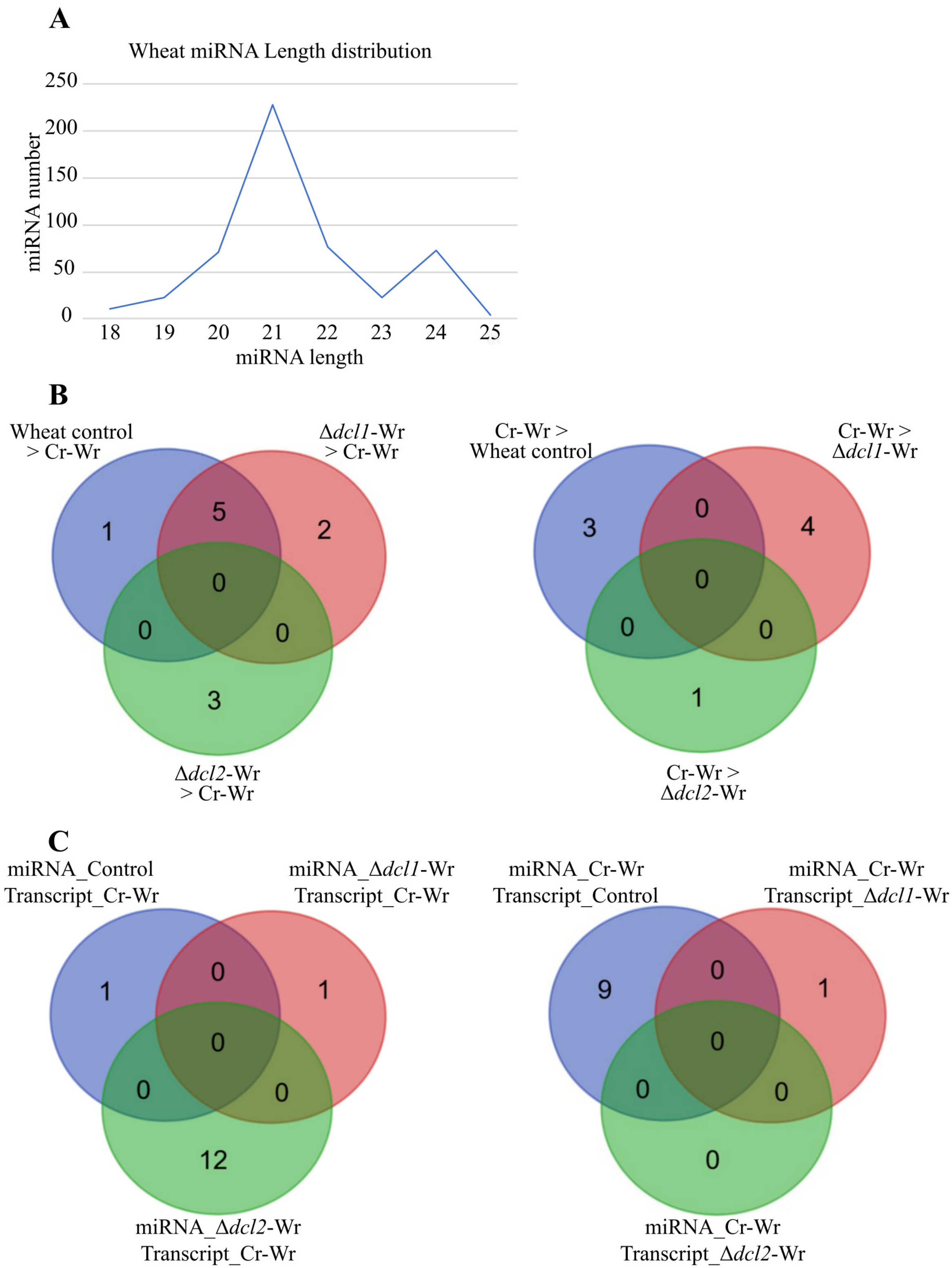

### Supplemental Figure 4

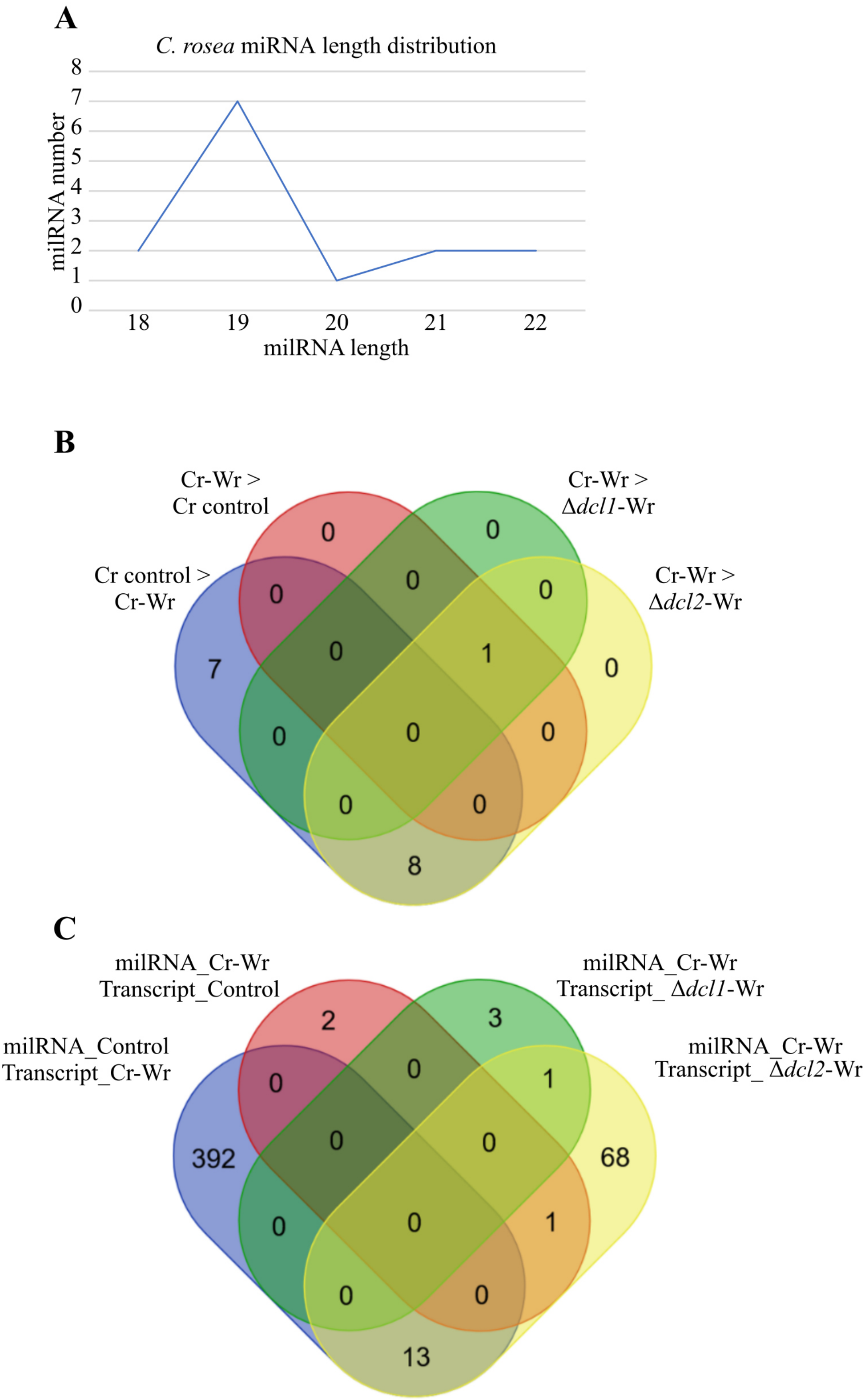
